# Structural variants underlie parallel adaptation following global invasion

**DOI:** 10.1101/2024.07.09.602765

**Authors:** Paul Battlay, Brandon T. Hendrickson, Jonas I. Mendez-Reneau, James S. Santangelo, Lucas J. Albano, Jonathan Wilson, Aude E. Caizergues, Nevada King, Adriana Puentes, Amelia Tudoran, Cyrille Violle, Francois Vasseur, Courtney M. Patterson, Michael Foster, Caitlyn Stamps, Simon G. Innes, Remi Allio, Fabio Angeoletto, Daniel N Anstett, Julia Anstett, Anna Bucharova, Mattheau S Comerford, Santiago David, Mohsen Falahati-Anbaran, William Godsoe, César González-Lagos, Pedro E. Gundel, Glen Ray Hood, Regina Karousou, Christian Lampei, Carlos Lara, Adrián Lázaro-Lobo, Deleon Leandro, Thomas JS Merritt, Nora Mitchell, Mitra Mohammadi Bazargani, Angela Moles, Maureen Murúa, Juraj Paule, Vera Pfeiffer, Joost A. M. Raeymaekers, Diana J. Rennison, Rodrigo S. Rios, Jennifer. K. Rowntree, Adam C. Schneider, Kaitlin Stack Whitney, Ítalo Tamburrino, Acer VanWallendael, Paul Y. Kim, Rob. W. Ness, Marc T. J. Johnson, Kathryn A. Hodgins, Nicholas J. Kooyers

## Abstract

Rapid adaptation during invasion has historically been considered limited and unpredictable. We leverage whole-genome sequencing of >2600 plants across six continents to investigate the relative roles of colonization history and adaptation during the worldwide invasion of *Trifolium repens*. Introduced populations contain high levels of genetic variation with independent colonization histories evident on different continents. Five large structural variants on three chromosomes exist as standing genetic variation within the native range, and exhibit strong signatures of parallel climate-associated adaptation across continents. Common gardens in the native and introduced ranges demonstrate that three structural variants exhibit patterns of selection consistent with local adaptation across each range. Our results provide strong evidence that rapid and parallel adaptation during invasion is caused by large-effect structural variants introduced throughout the world.

**Significance Statement:** Biological invasions occur over short timescales and introductions are often hypothesized to include limited genetic diversity, making the role of adaptation in invasion success controversial. We demonstrate that the invasion of a human-commensal species, *Trifolium repens*, likely stems from multiple, diverse introductions with significant evidence of climate-associated adaptation following introduction. The genetic basis of adaptation is most strongly linked to five chromosomal rearrangements that each span hundreds of genes – matching theoretical predictions that large-effect variants are key to the initial stages of adaptation to novel environments. Chromosomal rearrangements have remarkably parallel signatures of adaptation across different introductions despite initial colonization from different areas of Europe. Our study highlights the impact of globalization and rapid adaptation for the invasion success of human commensal species.

## Main Text

Invasive species threaten ecosystems, agriculture, human health, and culture. The cost of controlling the establishment, distribution, and spread of these species is immense (1–3), averaging US$26.8 billion per year globally. Yet, it is still not well understood why certain introduced species become invasive while others struggle to establish and spread. Despite substantial effort, research has identified few traits or processes that predict invasion across taxa (4–8). The role that introduction history and evolutionary processes like natural selection play in invasions have historically been neglected (9–11). However, recent literature stemming from large-scale experiments and the genomic revolution suggests that evolution occurs rapidly in invasive species and may be key to determining how invasions proceed - particularly in species that have been widely introduced and represent important components of ecosystems across the globe (12–16).

The role of adaptation in invasion success has been neglected because of the assumption that introductions are associated with severe bottlenecks that purge genetic variation and limit the potential to adapt (10, 17). However, many introductions and invasions do not fit this classic expectation, especially for human-associated species that are repeatedly introduced across the globe. Introduced populations may even have higher genetic diversity than native populations when admixture between genetically diverged populations occurs in the introduced range (18, 19). Natural selection and rapid adaptation have also been increasingly documented across invasive species (17). This paradigm shift leads to new questions that we address in this study. Specifically, how do introduction history and admixture interact to shape population structure during invasions? What is the genetic architecture of adaptation during invasions? And, does parallel adaptation occur across geographically-disparate introductions?

Theoretical population genetics predicts that rapid adaptation should be characterized by mutations of large effect (20). The limited literature on the genetics of adaptation during invasions largely support this hypothesis (15, 16), but genome-wide association studies (GWAS) and other quantitative genomic studies are biased toward larger effect variants (21). Many important traits for range expansion, such as an organism’s growth rate, size, and dispersal capability, have polygenic bases in diverse plant species. Adaptation via structural variants such as inversions may reconcile these observations. Inversions repress recombination, allowing clusters of co-adapted small-effect alleles to be inherited as a single segregating unit, (22), which have been linked to rapid adaptation (15, 23). However, inversions are difficult to link to fitness consequences across native and introduced ranges, as common gardens and reciprocal transplant experiments are logistically challenging and are rarely combined with large-scale genomic analyses.

Increasing globalization has resulted in repeated introduction of human commensal species across the world. Introduced species often encounter many of the same selection pressures throughout these ranges (e.g., altered climate regimes, release from herbivores, or loss of mutualists), thus some degree of parallel adaptation at the phenotypic and molecular scales might be expected (24). However, introduction history and demography may also have a significant impact on the distribution of adaptive and non-adaptive variation as the timing or order of the introduction of alleles may influence contemporary frequencies (i.e., priority effects (25)).

Admixture between genetically differentiated native populations in introduced areas can also create unique combinations of alleles that are only present in certain regions (26). Finally, if selected traits are polygenic, different loci underlying the same phenotype may be selected in each introduced region, reducing expected signatures of parallel evolution. Effectively parsing introduction history from adaptation requires genomic data from populations spanning similar climatic gradients in different areas of the world.

White clover (*Trifolium repens*) is an outcrossing legume native to Eurasia that has been introduced across the world as a forage and cover crop. Domestication occurred during the beginning of the second millennium in modern day Spain, and accessions spread across Western Europe and into the British Isles in the mid-1600s (27). Introduction to North America, South America, South Africa, Australia, Japan and China likely occurred as a result of European colonial activities, with establishment by at least the late 1800s in each region (28). Modern cultivars have been developed in North America, Australia, and New Zealand using germplasm sourced from across the world (29), and these cultivars were widely distributed in the second half of the twentieth century (30). Previous genetic studies in the native and various introduced ranges suggest that white clover is genetically diverse, with high effective population sizes in both native and introduced populations (31). Evolutionary ecology studies of a key defense polymorphism, cyanogenesis, have documented recurrent adaptive clines forming across climatic gradients and urban-rural gradients in native and introduced regions, suggesting that rapid adaptation has likely occurred following introduction (13, 24, 32–34).

Here we examine the interacting roles of introduction history and adaptation during the invasion of diverse climatic regions around the world. We utilized a low-coverage whole-genome sequence strategy to sequence the genomes of 2660 individuals from 13 populations spanning the native range of white clover, 39 populations from five introduced ranges, and twelve widely-used cultivars (Fig. 1; Data S1; Mean Coverage = 1.01x).

**Fig. 1:**
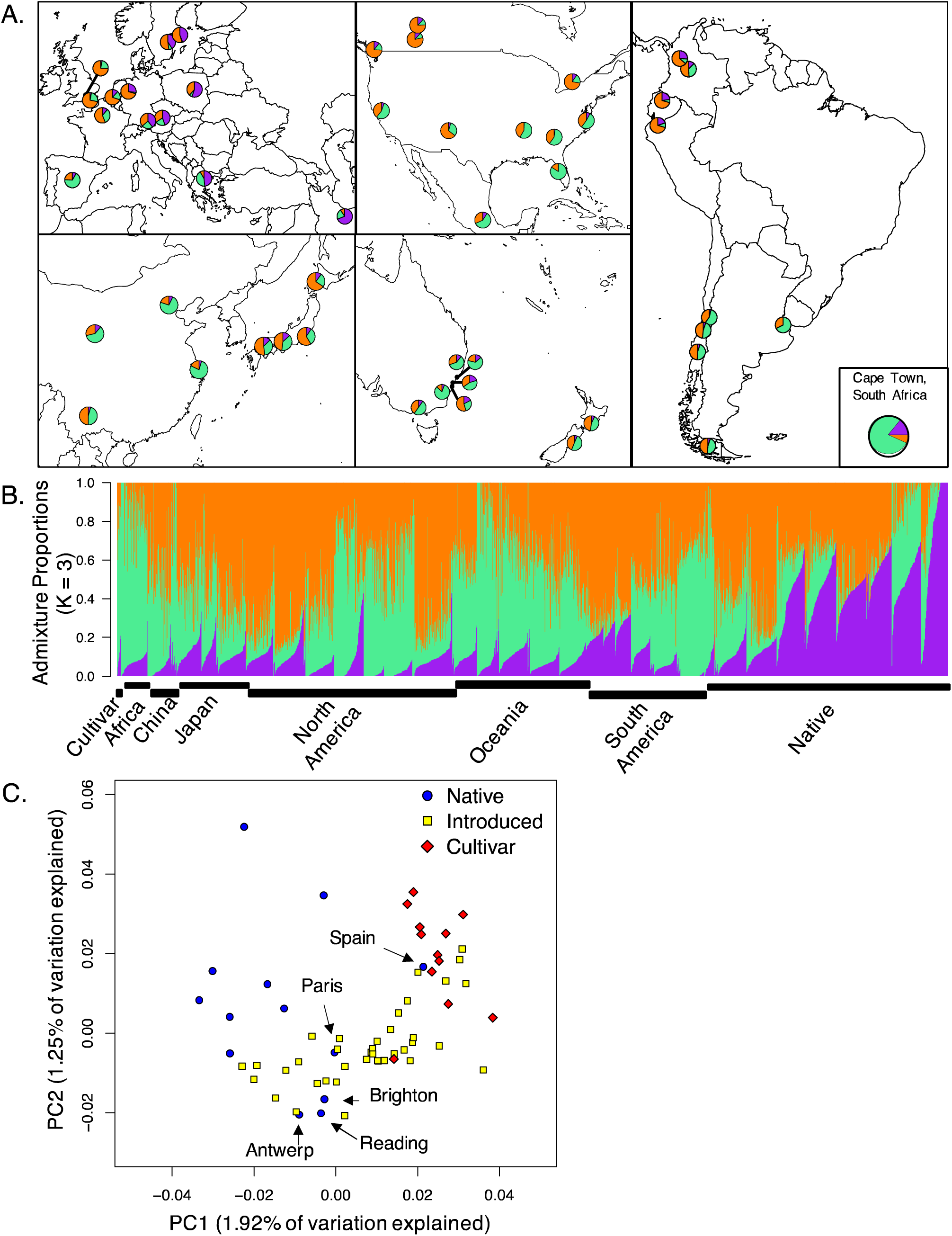
Population structure across worldwide populations of white clover. **A**. NGSadmix ancestries mapped across worldwide sampling. Each pie chart within map inserts reflects the average ancestries (K=3) from a given population. Note that Cape Town, South Africa, is the only population from Africa. **B.** Barplots depict ancestry output from the most likely K-value (K=3). Individuals are organized along the x-axis by population sorted by continent, longitude, and ancestry values. **C.** Principal component analysis visualizing population structure across ranges. Color and shape indicate geographic location of each population as shown in the legend. Population names and arrows refer to the native populations overlapping with introduced populations.

## Introduction History

White clover does not exhibit the classic signature of a population bottleneck in any introduced region. Genetic diversity is high in both the native and introduced ranges, with no clear difference in π (native π_Avg_ = 0.016, introduced π_Avg_ = 0.015; Welch’s ANOVA: F_1,21.465_ = 2.58, p = 0.12) or θ_w_(native θ_Avg_= 0.023, introduced θ_Avg_ = 0.019; Welch’s ANOVA: F_1,13.89_ = 1.23, p = 0.29). Despite this similarity, there is two-fold variation in genetic diversity among populations even within the same range (Fig. S1.; π_Range_ = 0.013 – 0.025), with Tampa FL, USA, having the lowest genetic diversity, and Edmonton AB, Canada, containing the highest diversity. Genome-wide Tajima’s D values are negative across both the native and introduced ranges, consistent with a recent population expansion (native D_Avg._ = -0.70, introduced D_Avg._ = -0.60). This pattern is expected given the recent worldwide introductions of *T. repens*. However, Tajima’s D does not differ between native and introduced ranges (Welch’s ANOVA: F_1,17.3_ = 0.30, p = 0.59).

Demographic modeling of effective population size (N_e_) during the recent past (1000 years) suggests that there is significant variation in N_e_ between populations with historic increases in N_e_ across most populations. However, this variation does not correspond to native vs. introduced populations and there are no signatures of recent bottlenecks or recent expansion across populations (Fig. S1). Together these results are consistent with the colonization of each introduced area involving repeated introductions of a high number of genetically diverse individuals.

We examined patterns of genetic differentiation between populations in the native and introduced ranges to better understand the relative independence of introduction events, different sources of introductions, and potential patterns of introgression between introduced ranges.

Consistent with high worldwide levels of genetic diversity and limited bottlenecks, genetic differentiation was low (Worldwide Average Weighted Pairwise F_ST_ = 0.027). Pairwise genetic differentiation was as strong within native and introduced regions as between regions (Fig. S2). There was a strong pattern of isolation-by-distance across the native range (Mantel’s r = 0.82, p = 0.001), with weaker patterns within introduced regions (North America: Mantel’s r = 0.18, p = 0.10; South America: Mantel’s r = 0.55, p = 0.002). These patterns are consistent with multiple and repeated introductions from the native region accompanied by additional gene flow across each introduced region. Pairwise differences between the native and introduced ranges were slightly elevated compared to pairwise differences between introduced regions, suggesting that introduced populations across the globe are more similar than expected. However, this varied by introduction region with higher differentiation between some introduced regions (South Africa-South America: F_ST_ = 0.04) than others (Japan-North America: F_ST_ = 0.02). Together these results suggest that there is significant admixture and/or shared colonization history between different introduced populations across the world.

To better parse population structure, we conducted admixture analyses using NGSadmix (35) using putatively neutral sites (i.e., four-fold degenerate). The most likely number of idealized populations was K=3 (36). Worldwide, all populations contained each of the three assumed ancestral gene pools (‘ancestries’) reflecting high levels of within-population variation. All three ancestries were strongly represented in different areas of the native range reflecting latitudinal and longitudinal patterns of isolation-by-distance (Fig. 1). The extreme east and west native populations (i.e., Tehran and Spain) represent distinct ancestries (purple and green, respectively), and a third ancestry increases with latitude (orange). Higher order K values (e.g., K=4,6, Fig. S3) further subdivide the native latitudinal gradient. Such population structure in the native ranges suggests that it should be possible to identify the strongest contributing populations to each introduction.

We compared ancestries of populations within the native and introduced ranges to detect the history of colonization and admixture. North American populations have ancestries most closely related to Spain in the south and France and Great Britain in the North. High-elevation populations in South America (Medellin, Bogota, Quito, etc.) and Japanese populations resemble high-latitude populations in North America (i.e. more orange ancestry). Lower elevation southern populations in South America, as well as Australian populations, New Zealand populations, Chinese populations, and South Africa, resemble southern populations in North America with similar ancestry coefficients to Spain (i.e. more green, Fig. 1). The similarities between different introduced areas likely reflect a shared introduction history as Western European nations brought white clover to these areas, but may also reflect post-introduction admixture between regions, or ecological sorting due to shared climate or biotic selection factors. For instance, Japanese and Chinese populations have very divergent ancestries that likely reflect differences in introduction history. However, parallel differences within continents, such as those observed in North and South America, may reflect contemporary admixture or ecological sorting across climatic gradients. Consistent with the ecological sorting, North America has a stronger pattern of isolation-by-environment than isolation-by-distance (Fig. S2).

To better determine the primary sources for each introduced region and determine whether the genomic variation in the introduced ranges is nested within the native range, we conducted a principal component analysis. Similarity in PC space closely corresponds to NGSadmix ancestries at K=3. There is differentiation among populations from native and introduced regions (Fig. 1C; PERMANOVA: F_1,49_ = 4.7, p = 0.039), with a limited number of native populations from Western Europe (Spain, Britain, France, Belgium) overlapping in PC space with the introduced populations. Similarity in PC space likely reflects colonization history, and it is notable that there is no clear clustering of different introduction regions. For instance, Canadian populations (Toronto, Calgary, Edmonton, and Vancouver) are located next to British, French, and Belgian populations, likely reflecting the introduction of white clover to these regions during French and UK colonization. Likewise, other North American populations are located midway between Spanish, French, and British populations, reflecting greater Spanish ancestry.

Introduction history does not explain all of the relationships within the PCA - introgression with modern agricultural cultivars could shape patterns of genome-wide population structure. To examine this hypothesis, we included 12 modern cultivars into our analyses. These cultivars were created in North America, Australia, and New Zealand with germplasm collected from populations in North America, France and New Zealand. Surprisingly, cultivars form a cluster distinct from introduced and native populations (PERMANOVA: F_2,60_ = 22.1, p = 0.001, Fig. 1C). With the exception of Grasslands Huia, cultivars are closely related to the Spanish populations and introduced populations from hot climates (Fig. S4). Thus, the cultivars do not necessarily reflect the regions where they were collected, but instead tend to have similar genetic compositions to one another. Nearly all of these cultivars were derived from field populations bred for resistance to drought and other environmental stressors. Conversely, Grasslands Huia, a New Zealand-derived cultivar, is closely related to other New Zealand wild populations. There is little doubt that admixture between cultivars and introduced populations occurs, but there remains significant differentiation from natural populations despite such introgression.

## Genomic Basis of Adaptation

Given the proliferation of white clover across the world in diverse habitats, an important question is: what role has adaptation played in the invasive spread of the species? Selection in introduced regions could favor different alleles that allow adaptation to novel conditions in the introduced range and/or that underlie traits that promote rapid invasion. We identify regions of the genome that show allele frequency differentiation between native and each of the five introduced regions using genome scans in 20 kbp windows (BayPass contrast (37); Fig. S5).

More differentiated regions of the genome (top 1% of windows) overlap between the native-introduced comparisons than would be expected by chance (hypergeometric test, p < 0.00001; Fig. 2), with the exception of comparisons with the Europe:Japan contrast, which did not show significant sharing with any other contrast (hypergeometric test, p = 0.16). These shared patterns of differentiation between introduced regions provide evidence for parallel selection pressures across introduced regions. However, no differentiated genomic windows are shared across all five introductions (Fig. 2), and few are shared across four regions (27 windows; 1.6% of windows that are an outlier for any contrast). Similar to the admixture analysis, North and South America share the most differentiated windows (128 windows, 29% of outlier windows). These results highlight parallel signatures of selection during range expansion across the species’ multiple introductions.

**Fig. 2:**
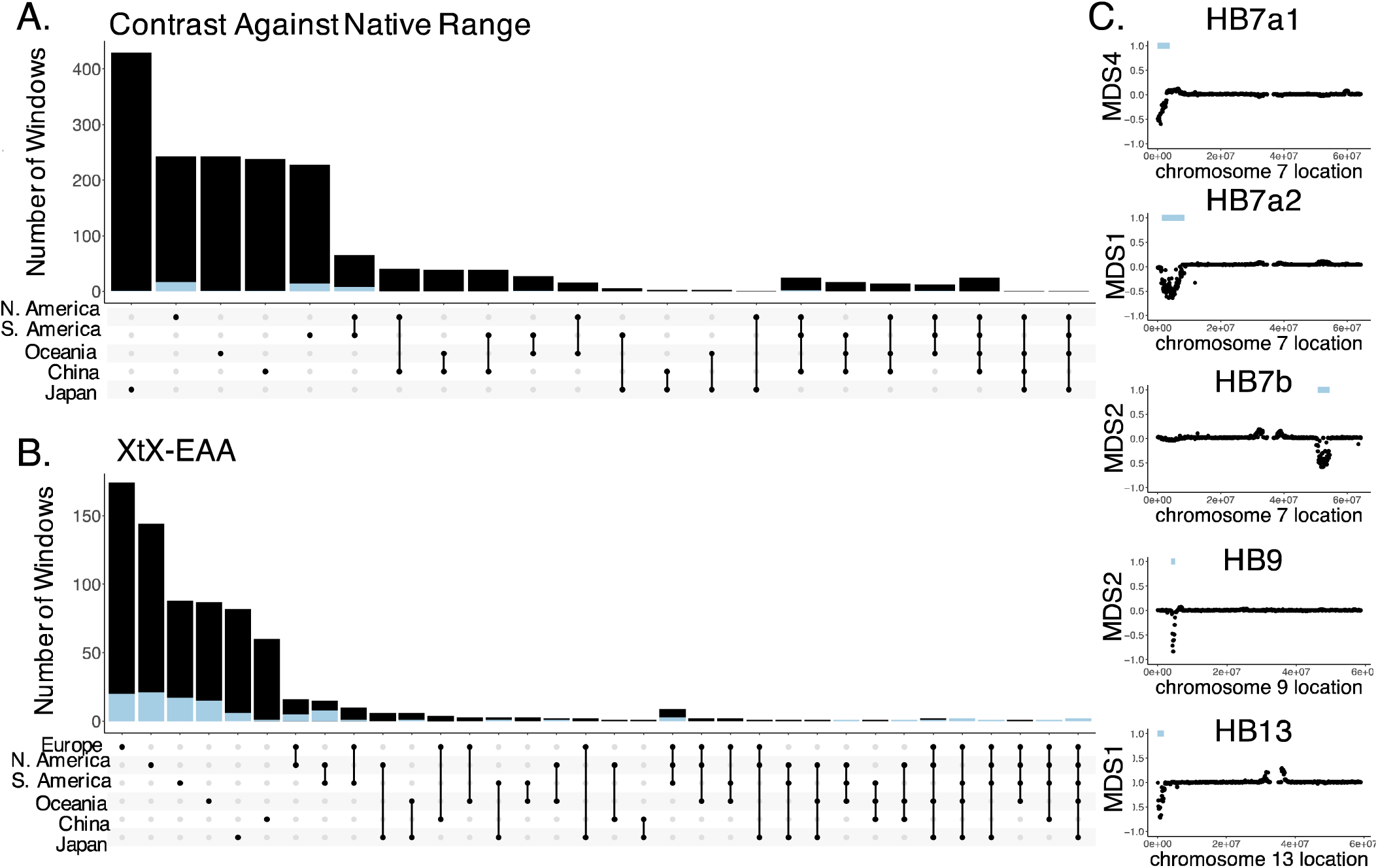
Signatures of structural variants are enriched for patterns of parallel selection across regions where white clover has been introduced. **A.** Upset plot depicting outlier windows for native:introduced region contrasts. **B.** Upset plot depicting windows corresponding to climate adaptation in each range (outliers for XtX statistic and correlations with at least one climate variable). Blue portions of bars in A and B correspond to genomic windows within haploblocks. **C.** Five haploblocks (putative structural variants indicated by blue bars above the region) identified as outliers on MDS axes summarizing local population structure along chromosomes.

Selection can also cause rapid adaptation to the environmental heterogeneity within each introduced range. We examined genomic regions underlying climatic adaptation in each introduced region by performing genome scans to identify 20 kbp windows enriched for sites showing both extreme population allele frequency differentiation (BayPass XtX (38)) and correlations with climate (39) (Fig. 3). In each range, between 15 and 52% of XtX outlier windows were also outliers for correlations with at least one of six minimally-correlated climate variables (XtX-EAA windows), indicating the importance of local climate adaptation following introduction (Fig. S6). Across the six ranges we observed signatures of genetic parallelism in climate adaptation, with all between-range overlaps of XtX-EAA climate adaptation windows significantly larger than would be expected by chance (hypergeometric test p-values < 0.013). Notably, there was some overlap between the windows identified in the contrast analysis and the XtX analyses (native range: 8.6%; introduced ranges: 7%). This pattern may be expected given that the sampled introduced ranges tend to have warmer climates than most of the native range (mean annual temperature: native = 10.3°C, introduced = 13.8°C, p = 0.006), and thus regions under climate-associated selection should be differentiated from the native range.

**Fig. 3.**
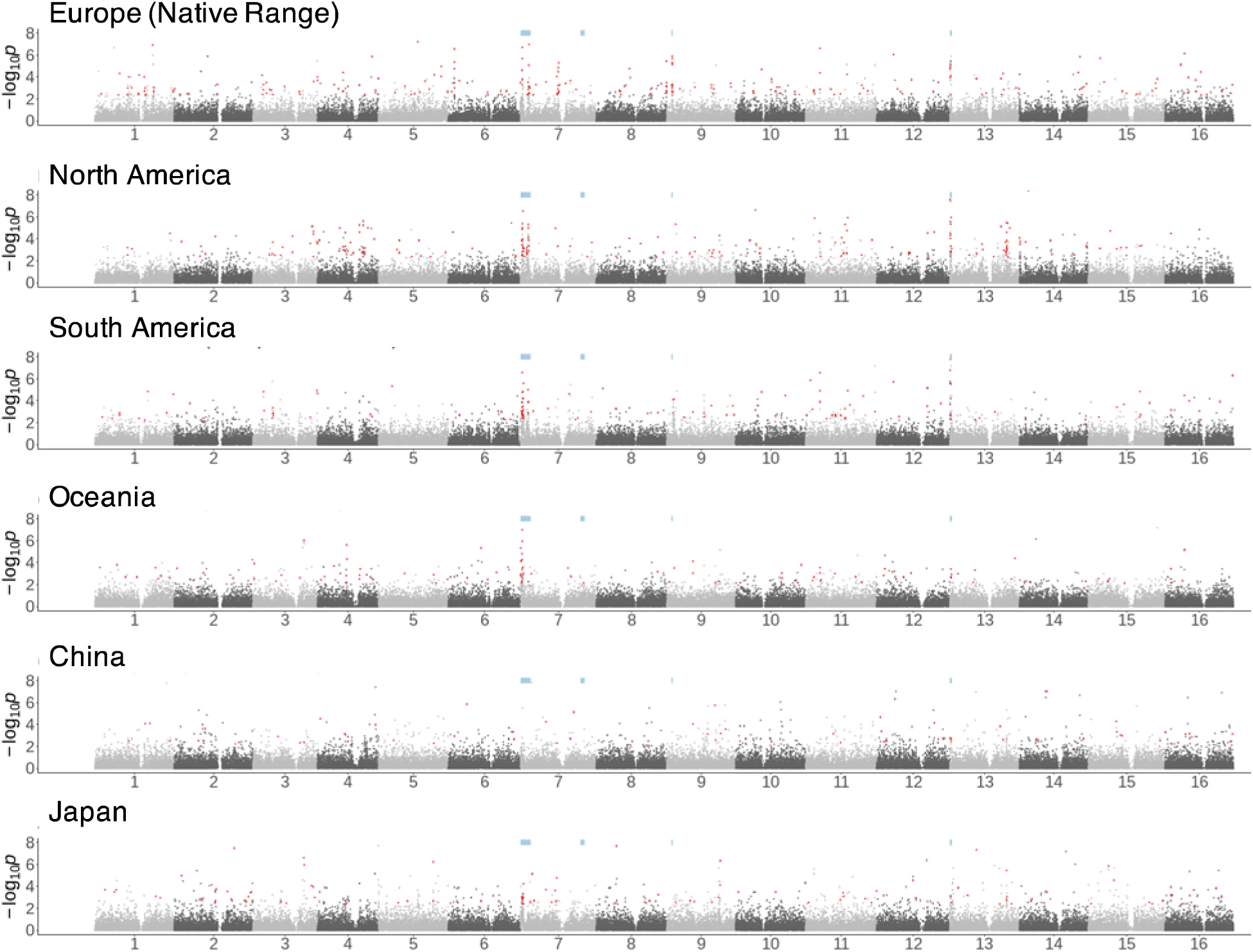
Haploblocks exhibit molecular signals of selection following introduction. *P*-values for enrichment of XtX (an F_ST_-like statistic that includes a correction for population structure) in 20 kbp windows across the genome using the weighted-Z analysis (WZA) within each white clover range. Red points indicate XtX-EAA outliers, windows that are in the 1% tail of WZA scores for XtX and correlation with at least one climate variable. Blue bars indicate haploblock locations.

The most notable peaks in each of the genome scans were several long stretches of differentiation (‘haploblocks’) on chromosomes 7, 9, and 13. Two partially-overlapping haploblocks on chromosome 7 (HB7a1 and HB7a2) and one haploblock on chromosome 13 (HB13) were shared among the Europe:North America and Europe:South America contrast comparisons (Fig. S5). Haploblocks HB7a1, and HB13 corresponded to dramatic signatures of climate adaptation in all ranges, while HB7a2 and HB9 showed strong signatures of climate adaptation in some ranges but not others. The breadth and synteny of these regions suggest that large structural variants may underlie convergent patterns of differentiation. We used a local principal component analysis of population-genomic data (15, 40, 41) to identify potential structural variants (i.e., inversions and translocations) across the genome. Potential variants contained stretches of windows with divergent population structure and exhibited higher levels of local nucleotide diversity within presumed heterozygous individuals compared to homozygous individuals. We identified signatures of five putative structural variants among 2,660 White Clover samples (Fig. 2, Fig. S7). Haploblocks HB7a1, HB7a2, HB7b, HB9 and HB13 were respectively 3.7, 7.1, 3.7, 1.2, and 1.8Mbp in size, and contained 591, 1014, 398, 152, and 227 genes. All haploblock reference and alternative alleles are found in nearly all of the populations in the native and all introduced ranges suggesting that haploblocks existed as standing genetic variation in the native range prior to introduction. However, allele frequencies differed between the introduced and native ranges for HB7a2 (t_49_ = -3.1, p =0.003), HB9 (t_49_ = 2.1, p =0.036) and HB13 (t_49_ = -2.2, p =0.03). Outlier windows for genetic differentiation (contrast) scans between the native range and North and South American invasions were significantly enriched for windows in haploblocks (Fig. S8; 14 and 12% of contrast outlier windows respectively; hypergeometric test p <= 9.67e-32). Haploblocks also have higher levels of within-range differentiation (XtX) than non-haploblock regions across every range except for China (Fig. S8), consistent with relatively strong selection on haploblocks to climatic variation within regions following introduction. These results suggest that large structural variants played an important role in range expansion following introduction.

## Characterization of Adaptive Structural Variants

To independently validate evidence for natural selection on the structural variants contributing to rapid adaptation following introduction, we conducted a transcontinental field experiment using diverse populations from the native and introduced ranges coupled with a genome-wide association study (GWAS). The experiment included four common gardens at low and high latitudes in both the native range (Uppsala, Sweden; Montpellier, France) and the introduced North American range (Lafayette LA USA; Mississauga ON, Canada). Each garden was planted with replicate plants from the same 96 natural populations; 47 populations collected along a latitudinal gradient in North America and 49 collected across Europe (33). Utilizing the same low-coverage whole-genome sequence approach as above, we genotyped 569 total individuals from this experiment for each of the five haploblocks. Frequencies of the reference and alternative haploblock alleles matched expectations from the worldwide population genomic dataset. We observed latitudinal clines in allele frequency in the predicted directions in North America for HB7a2, HB7b, and HB9 (Fig. 4). We did not expect latitudinal clines for HB13 or HB7a1 because allelic variation at these haploblocks does not differ between high and low latitude populations in the native and eastern North American ranges.

**Fig. 4.**
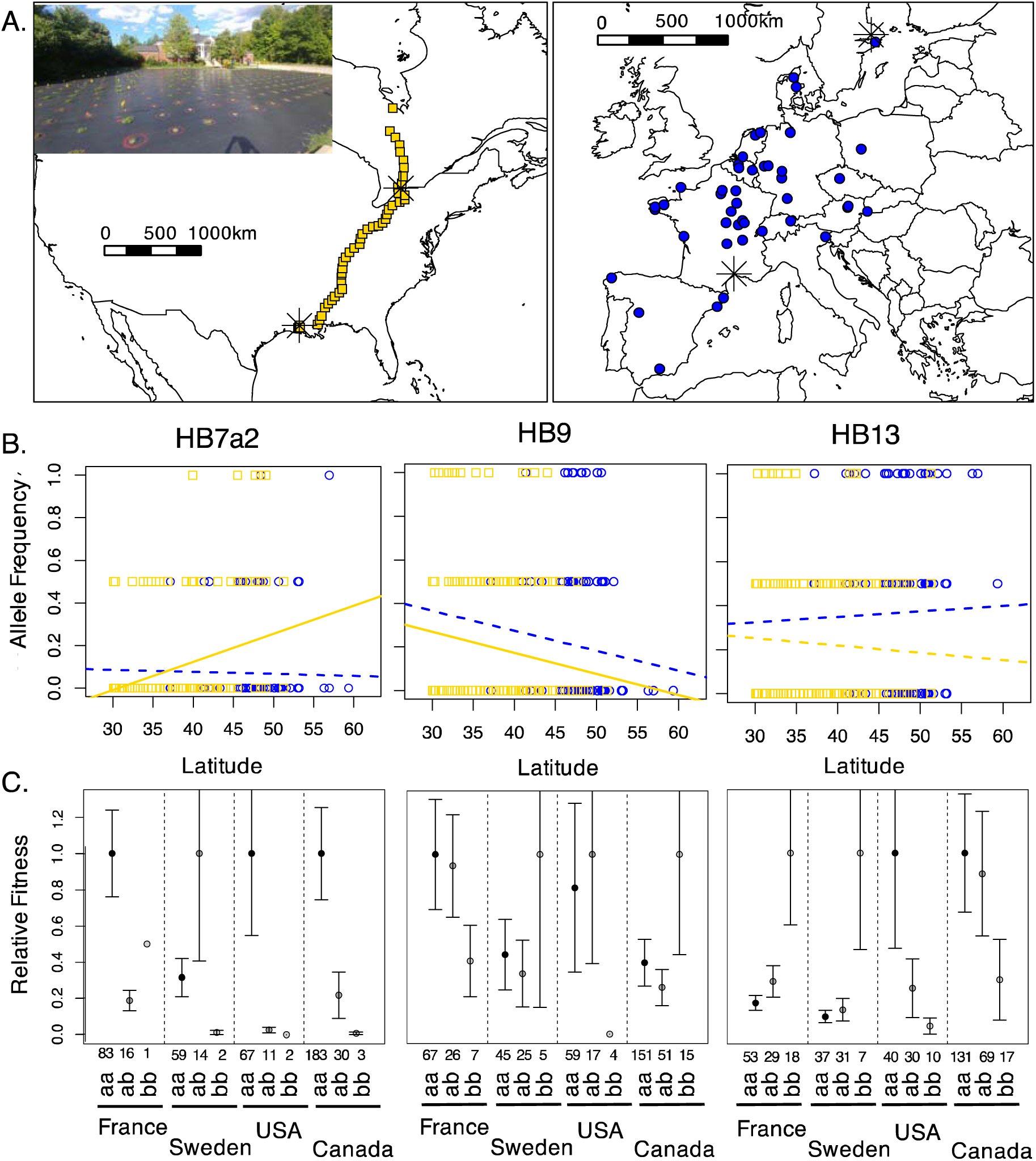
Impacts of haploblock variation on fitness across four field common gardens. **A.** Experimental design of the transcontinental field experiment. Points represent the 96 populations that were planted into each garden. Black asterisks are the locations of each garden. Insert picture is of the Mississauga ON, Canada garden. **B.** Alternative haploblock allele frequency for each individual from European (blue circles) populations or North American (gold circles) pooled across all four common gardens. Regression lines model allele frequency by latitude in Europe (blue) and North America (gold) with solid lines indicating statistically significant (P < 0.05) latitudinal clines and dash lines indicating non-significant regressions. **C**. Average relative fitness for each haploblock genotypes where the *a* allele represents the reference allele (56) and the *b* allele represents that alternative allele. Relative fitness was calculated from total seed mass and standardized by the genotype with the highest fitness within each garden. Error bars represent standard error around the mean.

We examined whether allelic variation at each haploblock influenced survival in the first year, growth rate, and relative fitness (i.e., total seed mass of an individual/mean total seed mass of all plants). There were significant garden x haploblock genotype effects on fitness consistent with haploblocks conferring local adaptation in the directions expected from the above genome scans (Fig. 4C, Table S1, Data S3). The strongest association was for HB13, where the alternative haploblock was strongly favored in the native gardens, but the reference haploblock was strongly favored in both North American gardens (ANOVA, Garden X Genotype: Χ^2^ = 9.6, p < 0.0001). Likewise, the HB9 alternative haploblock was marginally favored in the colder garden in both Europe and North America, while the reference haploblock was favored in the warmer gardens in both ranges (ANOVA, Garden X Genotype: Χ^2^ = 2.6, p = 0.05). Notably, the alternative allele for other haploblocks (HB7a1, HB7a2, HB7B) are at much lower frequencies which reduces our power to detect associations with fitness. Nevertheless, patterns at each haploblock still largely fit predictions established from allele frequencies. For instance, plants homozygous for the alternative HB7a2 allele had 92% greater survival in the first year in the Canadian common garden, but none of these homozygotes survived the first year in the Louisiana garden (ANOVA, Garden X Genotype: Χ^2^ = 7.5, p = 0.059; Table S1). These analyses provide experimental support that selection is operating on haploblocks, and the hundreds of genes contained within them, to drive rapid adaptation within introduced ranges.

We next evaluated which genes within each haploblock could be driving differences in fitness between gardens. We conducted separate GWAS within the native and introduced gardens. A number of loci in each haploblock were strongly associated with the ability to flower or total seed mass (Fig. 5; Table S1). Most hits were observed in the North American gardens due to sample size differences between gardens. The number of hits exceeded the genome-wide expectation for each haploblock for at least one fitness measure (Fig. S9). Hits were located within 10 kbp of annotated genes, but only two hits fell directly within the coding sequence of a predicted gene. The number and location of fitness-associated SNPs within putative structural variants suggests that there are multiple genomic regions under selection within each haploblock and that differential expression is likely the important driver of adaptive phenotypic differences.

**Fig. 5.**
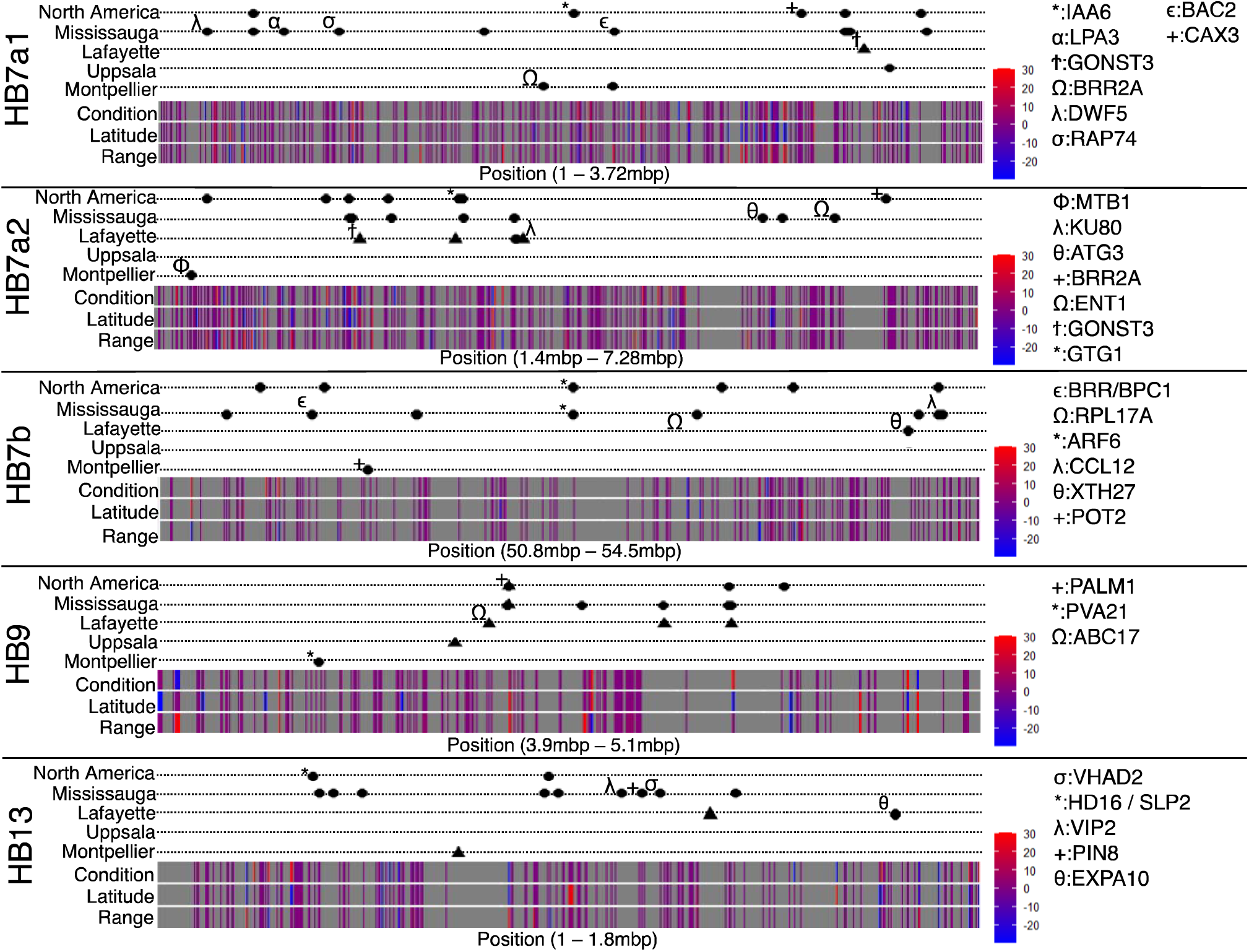
Associations with fitness and differential expression within haploblocks. GWAS hits for survival to flowering (triangles) and total seed mass (circles) are plotted above expression ribbons. Each dotted line corresponds to a GWAS analysis within a particular garden (Mississauga, Lafayette, Uppsala, or Montpellier) or a GWAS analysis that merged the North American Gardens (North America). Hits that either fall within a coding sequence or are within 10 kbp of a known homologue are marked with a Greek or mathematical symbol as shown in the legend to the right of each graph. Differential expression ribbons visualize expression patterns for each gene within comparisons between condition (drought vs. well-watered), latitude (low vs. high) and range (native vs. North American). Heatmaps for differential expression for each gene range from down-regulated (blue) to up-regulated (red). Full descriptions of Fitness GWAS hits and differentially expressed genes can be found in Data S4 and Data S5, respectively.

The genes nearby fitness-associated SNPs within the haploblocks correspond to stress resistance, defense and flowering, matching expectations from a GO analysis of these regions (Data S2). Of the multiple fitness-associated SNPs within the HB7a1 haploblock, one of the most prominent was found downstream of *IAA6* (indole-3-acetic acid 6; p = 1.32e-5, β = -0.97), a key regulator of auxin responses, phototaxis and development in Arabidopsis (42). The two GWAS hits underlying survival to flowering on HB7a2 were associated with *MT1B* (p = 2.18e-7, β = 1.24) and *GTG1* (p = 2.10e-6, β = -1.187). *MT1B* acts as a reactive oxygen species (ROS) scavenger in rice, protecting plants from water stress and metal toxicity (43). *GTG1* encodes a conserved membrane protein that plays a role in root growth and light responses in Arabidopsis (44). Two GWAS hits underlying survival to flowering on HB7b were within the coding sequence of *ARF6* (p = 2.24e-6, β = -1.73); *ARF6* is characterized as a transcription factor involved in flower maturation in Arabidopsis (45). Notably, multiple genes associated with photoperiodic control and flowering in other species are associated with survival to flowering within the HB13 haploblock including hits downstream of *Hd16* (p = 3.61e-5, β = -1.03; (*46*) and *SLP2* (p = 3.61e-5, β = -1.03; (*47*). Identification of these genes suggests that each haploblock contains ecologically-important variation that is likely central for rapid adaptation following invasion and provides specific targets for downstream functional analysis.

We further validated the fitness-associated SNPs within the haploblocks above with a manipulative RNAseq experiment conducted in growth chambers. We evaluated genome-wide differential expression between high- and low-latitude populations of white clover from the native and invasive range both in dry down and well-watered conditions. The water availability treatment was selected because the differential mortality between common gardens was hypothesized to be associated with drought. While elevated, differentially expressed genes were not overrepresented within haploblocks compared to the rest of the genome for any comparison (treatment, range or latitude; Fig. S10) and the magnitude of expression changes for differentially expressed genes within haploblocks was similar to the rest of the genome (Fig.

S10). However, a high percentage of hits in the fitness GWAS above were differentially expressed in a least one comparison (Survival to Flowering GWAS hits: 38.0%, 68 of 179 genes; Total Seed Mass GWAS hits: 30.6%, 11 of 36 genes). This group of genes was relatively uniformly distributed across the different haploblocks, i.e. Survival to Flowering GWAS hits represented 25-48% of differentially expressed genes across haploblocks. This group included twelve genes with clear orthologs (Survival to Flower: *ARC11, ATG3, CCL12, ENT1, EXPA10, IAA6, MT1B, PIN8, RAP74, RPL19, SLP2*; Total Seed Mass: *GONST3*) that were associated with fitness in GWAS analyses and had differential expression across drought treatment, range and latitude (ANOVA, Treatment x Range x Latitude: p_adj_ < 0.0001; Data S5). Several of these genes (including *IAA6*, *MT1B*, *ATG3*, *EXPA10*, *RPL19*) have been associated with drought and oxidative stress in other species (43, 48–51). In sum, we find that the same genes identified in the fitness GWAS have different constitutive patterns of expression in populations from different ranges and latitudes. This result is consistent with cis-regulatory changes underlying rapid adaptation following introduction, but does not exclude the possibility that haploblocks also include ecologically important variation in protein-coding regions.

## Conclusions

We demonstrate that the invasion of white clover throughout the world has been achieved through a complex pattern of global colonization accompanied by rapid adaptation. While worldwide patterns of population structure reflect some aspect of colonization history and independent introduction events, our demographic analyses are consistent with white clover undergoing multiple, repeated introductions, followed by admixture among diverse ancestral haplotypes. This complex introduction history has maintained substantial genetic diversity and high effective population sizes in introduced populations, including the formation of novel genetic variation.

In addition to complex demography, we observe striking signatures of climate-related selection in introduced areas around the world. While there are clearly many regions across the genome with evidence of selection, the strongest and most parallel signatures of adaptation come from just a few haploblocks that also exhibit classic genomic signatures of structural rearrangements (i.e. inversions and translocations). Allelic variation within haploblocks is strongly associated with differences in relative fitness between common gardens in the native and introduced North American range, demonstrating that haploblocks underlie patterns of local adaptation that have evolved in the last 400 years. Variation within these haploblocks suggests that the molecular basis of these differences lies in differential expression of key genes involved in the developmental timing, stress tolerance, and defense.

Our results demonstrate the power and importance of rapid adaptation during an invasion (14, 17, 52). We identify large-effect structural variants as driving rapid adaptation to novel environments, which match theoretical expectations from the modern synthesis (20, 53) as well as earlier empirical studies on individual inversions in *Drosophila* (54). However, unlike theoretical models that rely on *de novo* mutation, each haploblock that we identify exists as standing genetic variation in the native range, and repeated introductions facilitated the movements of such variation to different regions around the globe. Lag periods preceding rapid expansion during invasions may not only be an opportunity for demographic increase and sorting, but also an opportunity for additional input of standing variation from the native or other introduced ranges. We suggest that selection and adaptation are likely the norm for human-commensal species that are commonly introduced, which contributes to invasion success throughout the world.

## Materials and Methods

*Trifolium repens* L. (white clover) is an outcrossing herbaceous perennial that has a cosmopolitan distribution across regions with temperate climates. White clover is native across Eurasia and was subsequently domesticated in Spain between 1000-1200AD as a forage and nitrogen-fixing rotation crop. Introductions across the world coincided with European colonization. Breeding of modern white clover cultivars has occurred in several regions where white clover is an important crop including New Zealand, North America, Australia, and China.

### Population genomics dataset

Our dataset includes low coverage whole genome sequences from 2616 samples collected from 50 different cities spanning the native range in Eurasia (12 cities) as well as introductions to North America (11 cities), South America (10 cities), Japan (four cities), China (four cities), Oceania (eight cities), and Africa (one city).These samples were collected as part of the Global Urban Evolution Project from 2016-19 (34). Each city was treated as a single population and sample sizes for each population ranged from 5-120. This heterogeneity in sample size was intentional as we wanted to include a number of cities with high sampling for better estimates of site frequency spectra and population-genomic statistics (31 cities; Ave. = 80.74, Std. = 17.7 individuals; 52). We then added additional cities with lower sampling that we deemed as important areas for understanding colonization history (19 cities; Ave. = 5.95, Std. = 0.23 individuals). Additionally, we sequenced 32 samples collected from four cities in Spain (A Corona, Granada, Salamanca, San Sebastian; 33) as well as 12 popular cultivars bred in the U.S. (Durana, Patriot, Renovation, Merit, Pilgrim, LA-S1, CA Ladino), Australia (Irrigation), and New Zealand (Crau, Grassland Huia, Grasslands Pitua). Details on library construction and sequencing for new samples is described in the Supplemental Methods. Environmental data for each sampling location was extracted from BIOCLIM using the *raster* v3.6-26 package in R.

### Analysis of demography and worldwide population structure

Sequences were processed using a common pipeline (https://github.com/James-S-Santangelo/glue_dnaSeqQC) and aligned to a chromosome-level genome assembly (56). For demographic analysis, we extracted 4-fold degenerate sites using the Degeneracy Pipeline (https://github.com/tvkent/Degeneracy) and utilized all sites for genome scans. We assessed population genomic diversity, differentiation, and structure using genotype likelihoods in ANGSD v0.929 (57). To examine genetic diversity within each population, we first calculated genotype likelihoods and site allele frequency likelihoods (SAF) for each population independently using only 4-fold degenerate sites (*-GL 1 -doMaf 2 -doCounts 1 -dumpCounts 2 - baq 2 -minQ 20 -minMapQ 30 -doSaf 1 -sites 4fold.sites*). 1DSFS were used to calculate thetas (θ_w_ and θ_π_) using *realSFS saf2theta* and *thetaStat do_stat*, while 2DSFS were used to estimate differentiation using Hudson’s Fst (*realSFS fst index -whichFst 1*). Average number of SNPs per population for these analyses was 10,784,068 (SD 865,692).

We identified signatures of bottlenecks by comparing genetic diversity statistics and Tajima’s D between native and each introduced region. We estimated Ne through time using a coalescent framework implemented in EPOS (58), focusing on population contractions in the last 1000 years as signatures of bottlenecks. We investigated patterns of genetic differentiation within and among populations across the native and introduced ranges by calculating pairwise weighted and unweighted Fst values using ANGSD (59, 60). Isolation by distance and isolation by environment in native and introduced ranges were assessed via Mantel tests using the *mantel*() function within the *vegan* library (61) with Haversine geographic distance matrices via *distm*() function within the *geodist* library and climatic distance matrixes using the *dist()* function in the *vegan* library. We examined worldwide population structure and individual ancestry using NGSadmix (35). NGSadmix runs included 3-8 replicates of K=1-8 using 10,000 iterations per replicate (*-maxIter*). To determine the most likely number of clusters, we examined standard deviations in likelihoods at each K and used the method of Evanno et al. (36) to identify the most likely number of ancestral clusters and the uppermost level of population structure. To better dissect introduction history, we examined patterns of nested population structure using principal component analysis. We used *PCAngsd* (62) to generate a variance-covariance matrix using genotype likelihoods and estimated allele frequencies (*pcangsd.py*), and then extracted the eigenvectors (i.e. the principal components) of the covariance matrix using *eigen()* function in R. To examine potential clustering within the PCA, we conducted PERMANOVA using the *adonis2*() function within the *vegan* library (61). Difference in number of samples or sequencing coverage have limited impact on our inferences of population structure (Fig. S11).

### Genome scans for signatures of selection

We identified regions of the genome under selection using two separate approaches. First, we contrasted allele frequencies in native range with those in each invasive range. Second, we looked for relationships between allele frequency and climate within each individual range as evidence of local climate adaptation. Genotype likelihoods were calculated in ANGSD (*-GL 1 - doGlf 2 -doMajorMinor 4 -doMaf 2 -baq 2 -minQ 20 -minMapQ 30 -SNP_pval 1e-6 -minMaf 0.05*) in each range (Europe, North America, South America, Oceania, China and Japan) for climate adaptation scans or pair of ranges for contrast scans. We then estimated population allele frequencies for these sites in each population individually using ANGSD (*-GL 1 -doGlf 2 -doMajorMinor 4 -doMaf 2 -doCounts 1 -baq 2 -minQ 20 -minMapQ 30 -minMaf 0*). Allele frequencies for sites were only used if they were callable for all populations in a particular scan (14.7M-22.7M sites/range).

We used the BayPass contrast statistic (37) to summarize allele frequency differentiation at each site between European populations and populations from an invasive range. Enrichment of contrast outliers was calculated for non-overlapping 20 kbp windows using the weighted-Z analysis (WZA, 60) and outlier windows were defined as the 1% tail of the distribution of WZA window scores.

We tested for genomic regions with greater differentiation than expected by chance within each native range while accounting for genome wide population structure using the BayPass core model. For these genome scans, we generated population covariance omega matrices for each range in BayPass v2.2 (37, 38) using 10,000 sites sampled from outside annotated genes. We then ran the BayPass core model to quantify allele frequency divergence between populations within each range while accounting for population structure using the omega matrix (XtX). Next, correlations between population allele frequencies in each range and six minimally correlated bioclimatic variables (BIO1, BIO2, BIO8, BIO12, BIO15 and BIO19 from the WorldClim dataset, 39) were quantified using the absolute value of Kendall’s Tau. In each range, we used WZA to identify non-overlapping 20 kbp windows that were enriched for outliers for the XtX statistic and correlations with each bioclimatic variable. Outlier windows for each statistic were defined as the 1% tail of the distribution of WZA window scores. Outlier windows that overlapped between genome scans were identified, and their enrichment relative to a hypergeometric distribution was tested in R.

### Haploblock identification

We identified population-genomic signatures of haploblocks using local principal component analysis, modifying the method described by Li and Ralph (40) to utilize covariance matrices from PCAngsd v1.10 (62), which were calculated in 100 kbp windows from beagle files generated in ANGSD v0.929 (5) (*-GL 2 -doMajorMinor 1 -doCounts 1 -doGLF 2 -SNP_pval 1e- 6 -doMaf 2 -doGeno -1 -doPost 1 -minMapQ 30 -minQ 20 -trim 5 -minMaf 0.05 -minInd 665 -geno_minDepth 2 -setMinDepthInd 2 -uniqueOnly 1*). Local population structure along each chromosome was analyzed on five MDS axes and outliers were identified from the 5% corners of each pair of MDS axes. Heterozygosity was also calculated for each sample in each candidate region using ANGSD (*-dosaf 1 -minMapQ 30 -minQ 20 -trim 5 -GL 2*) and realSFS v0.929 (64) (*-fold 1*). After filtering out samples with less than 0.4X coverage, putative inversions were identified by the presence of three clusters of samples along a single principal component axis, indicative of two homozygous and one heterozygous inversion genotype, as well as by the presence of elevated heterozygosity in the region among heterozygotes. We defined elevated heterozygosity as when the standard error of the mean (SEM) for each homozygote class did not overlap the SEM for the heterozygote class. We validated haploblock genotypes by performing LD scans with ngsLD v1.2.0 (*--min_maf 0.05 --max_kb_dist 0*) on 5000 randomly sampled sites from each chromosome containing a haploblock. For each haploblock, LD scans were run on a set of samples homozygous for the more common haploblock allele, as well as a random set of samples of the same size. To identify GO terms enriched in haploblocks, the *topGO* library (65) was used with Fisher’s exact test, the ‘weight01’ algorithm, and a *p*-value < 0.05 to assess significance.

### Associations between haploblocks and fitness

We examined patterns of local adaptation and the genomics of adaptation using four common gardens located in the southern and northern region of the native range (Montpellier, France and Uppsala, Sweden respectfully) and the southern and northern regions within the North American introduced range (Lafayette, USA and Mississauga, Canada respectfully). Common gardens were conducted for two years at each site, 2020-21 in North American gardens and 2021-22 in European gardens. Seedlings from the same lines were planted in each garden. Seeds were collected from 46 white clover populations spanning a 27° latitudinal gradient in Europe and from 47 additional populations spanning a 21° latitudinal gradient in North America. Seeds were grown for a single refresher generation and outcrossed via hand-pollination within each population. We established 4-6 outbred lines per population before randomizing and planting directly into the natural soil of a cultivated lawn at each site. Survival was surveyed and mature fruit were collected weekly. We report two measures of fitness: “Survival to Flowering” is a binary variable that indicates whether a plant was able to flower during the two-year experiment and represents both viability and ability to mate. “Total seed mass” reflects both viability and fecundity as plants that did not produce any seeds had no seed mass.

We generated low coverage whole genome sequences for 569 samples across the four gardens using the same library construction and bioinformatics pipelines as above. We estimated haploblock genotypes by performing local PCAs on each previously-identified haploblock region including all lcWGS samples. The first two principal components of genetic variation across GLUE and common garden samples for each haploblock region were visualized and used to assign common garden samples to genotype clusters.

We assessed whether haploblocks were associated with adaptation following introduction via a three-pronged approach. We first validated our GLUE dataset by using linear models (*lm()*) to identify associations between haploblock genotype and latitude of collection site. We then qualitatively compared whether clines in the native and introduced region in these gardens matched the GLUE dataset. Second, we examined how haploblock variation impacts fitness across gardens. We modeled survival to flower and total seed mass in separate univariate generalized linear models with Garden, Genotype, and Garden:Genotype interaction as factors. GLMs were implemented using *glm()* and statistical significance of each factor was assessed using *Anova()* with Type III sum of squares in the *car* library (66). Survival to flower was modeled with a binomial error distribution and a logit link. Total seed mass was log(+1) transformed and modeled with a gaussian distribution and identity link. Finally, we calculated relative fitness from total seed mass data for each haploblock to better understand the strength of selection acting on each haploblock within each garden. Relative fitness for each haploblock was calculated by dividing each individual value for total seed mass by the average value of total seed mass for the genotype with the highest fitness in the garden.

To identify the genes and phenotypes potentially under selection, we conducted genome-wide association studies (GWAS) with a genotype-likelihood framework implemented in ANGSD. We conducted independent GWAS for the two fitness traits in each garden. Then, we pooled European gardens and North American Gardens and conducted GWAS for each trait in each pooled sample. Genotype likelihoods were estimated for each range (174 individuals from the native European range and 395 individuals from the introduced North American range) in ANGSD (*-GL 1 -minMaf 0.05 -minMapQ 30 -minQ 20*). GWAS employed a hybrid model (*-doAsso 5*) that first uses a score statistic to evaluate the joint maximum likelihood estimate between a trait and an observed marker (67). If the chi-square test falls below a particular threshold (*-hybridThres 0.05*), a latent genotype model with an expectation-maximization (EM) algorithm is fit (68). We controlled for population structure by adding the first 20 principal components as covariates. Principal components were generated in *PCAngsd* as above. In GWAS for each combined range, we also added garden as a covariate. To account for multiple tests, we used a conservative Bonferroni correction. We used permutation analyses to determine whether the number of fitness GWAS hits exceeded expectations from the rest of the genome (see supplement).

### Differential Expression Analysis

We performed a manipulative experiment to examine variation in expression between white clover populations in native and introduced range under dry down and well-watered conditions. We selected 1-3 maternal lines from each of 3-4 populations from low and high latitudes in the European and North American ranges (14 total populations; 47 total samples). Plants were grown for six weeks to accumulate above- and below-ground biomass. At six weeks, all pots were saturated with water by bottom-watering. Plants in the control (well-watered) flats received periodic watering according to our standard greenhouse protocol. Plants in the dry down treatment did not receive additional water. Each day, we assessed soil moisture in each pot using a SMT150T soil moisture meter (Dynamax; Houston, TX). Leaf tissue from two healthy adult leaves was flash frozen in liquid nitrogen 10 days after the dry down treatment began from plants in both the well-watered and control treatment. Library construction, sequencing, and bioinformatics details in Supplemental Methods. We used DESeq2 (69) to test for differences in transcript abundance between dry down and well-watered treatment groups, between the North American and European ranges, and between high- and low-latitude populations. We used two different models to examine differential patterns of gene expression across treatments, range and latitude. The first included all interactions (treatment*range*latitude). The second set of models were univariate models examining differential expression across treatment, range and latitude separately. Genes were categorized as differentially expressed if FDR was < 0.1. We evaluated whether transcribed genes located within haploblocks were more or less differentially expressed than in other regions of the genome by resampling across the genome. Briefly, the same number of genes found within each haploblock were randomly sampled across the genome 10,000 times while preserving synteny. The number of genes with an FDR < 0.1 for each of the 10,000 sampled haploblocks was summed and the average log2Foldchange was calculated, which were then used to create a null distribution for each haploblock region of the expected number of differentially expressed genes and their relative log2Foldchange.

## Acknowledgments

This work would not be possible without white clover collections by 287 fellow scientists in the GLUE network. Support for field work was provided by Andre Daugereaux, the UL-Ecology Center, and numerous field and lab assistants in each common garden. The Louisiana Optical Network Infrastructure provided computational support. The use of genetic data originating from Ecuador was approved by the Ecuadorian Ministerio del Ambiente (access permit MAE-DNB-2018-0106 & transfer permit ATM-CM-2018-0106-001– 2019). Funding was provided by an NSERC CGS Doctoral Award to LJA; NSF grants OIA-1920858 and DBI-2244712 to N. Kooyers; CNRS-University of Toronto Ph.D. Student Travel Grant to MTJJ and CV; NSERC Discovery Grant, Canada Research Chair, and NSERC Steacie Fellowship to M. Johnson; and an ARC DP220102362 and HFSP RGP0001 grant to K. Hodgins.

## Author contributions

Conceptualization and Methodology: PB, BTH, JIMR, JSS, LJA, PYK, RWN, MTJJ, KAH, NJK

Resources: JSS, LJA, SGI, FA, DNA, JA, AB, MSC, SD, MFA, WG, CGL, PEG, GRH, RK, CL, CL, ALL, DL, TJSM, MMB, AM, MM, JP, VP, JAMR, DJR, RSR, JKR, ACS, KSW, IT, AVW, MTJJ, NJK

Investigation: LJA, NK, AP, AT, CV, FV, CS, CMP, PK, MTJJ, NJK

Dataset Curation: PB, BTH, JIMR, JIMR, NJK

Formal Analysis: PB, BTH, JIMR, JSS, JW, AEC, MF, NJK

Funding acquisition: LJA, CV, MTJJ, KAH, NJK

Project administration: MTJJ, KAH, NJK

Supervision: MTJJ, KAH, NJK

Writing – original draft: PB, BTH, MTJJ, KAH, NJK

Writing – review & editing: All

## Data and materials availability

Low coverage whole genome sequences (.bam files) for all accessions can be found as .bam files in the NCBI SRA database (removed before publication). Metadata and fitness data from the four-way common garden study can be found on Dryad (removed before publication). Raw fastq files from RNAseq expression experiment can be found in the NCBI SRA database (removed before publication). All code from this manuscript is linked to a github repository (github.com/pbattlay/glue-invasions).

## Supplementary Materials

Supplementary Materials and Methods Figs. S1 to S9 Table S1 Data S1 to S5

