## Supplemental Material for "Structural variants underlie parallel adaptation following global invasion"

**The PDF file includes:**

Supplemental Materials and Methods
Figs. S1 to S11
Table S1

**Other Supplementary Materials for this manuscript include the following:**

Data S1 to S5

**Supplemental Materials and Methods**

In this study, we combine a population genomic analysis across worldwide populations of white clover (*Trifolium repens* L.) with genome-wide association analysis of fitness within native and introduced populations and RNAseq analysis in a manipulative experiment. For each of these experiments, we describe the dataset, the experimental design, and the respective analyses presented within the main text.

Study system

*Trifolium repens* L. (white clover) is an outcrossing herbaceous perennial that has a cosmopolitan distribution across regions with temperate climates. White clover is an allopolyploid (4n=32) with disomic inheritance derived from two western European progenitors, *T. occidentale* and *T. pallecens* approximately 15,000 - 28,000 years ago (1). Following hybridization, *T. repens* expanded its range across all of Europe and Middle East. Here we consider all of Eurasia as the native range. White clover was subsequently domesticated in Spain between 1000-1200AD as a forage and rotational crop, specifically for its nitrogen-fixing capacity and stoloniferous vegetative growth (2). Domesticated white clover spread rapidly around Western Europe and obtained the common name 'dutch clover' due to extensive proliferation in the 1500's throughout present day the Netherlands. Domesticated versions of white clover were documented in the UK in the early 1600s. Subsequent spread around the world occurred rapidly with European colonization, as white clover (along with red clover) represented the main mechanism of fertilization in many crop systems (3). Timing of introduction to North America, South America, South Africa, Australia, Japan and China likely occurred soon after colonization. For instance, in North America, journal entries by Benjamin Franklin and observations by botanist Petr Kalm suggest that white clover existed as a feral commensal species across eastern North America by 1749 (3). First records in other regions are less well documented, but white clover likely established by the late 1800s in each region.

While the nitrogen-fixing capacity of white clover was less necessary after improvements to chemical fertilizer production in the early 1900s, breeding of modern white clover cultivars has occurred in several regions where white clover is still deemed an important forage or cover crop including New Zealand, North America, Australia and to a more limited degree, China. In the agricultural literature, white clover is largely separated into three different types based on leaf size and growth habit (4, 5). Ladino clover has large leaves, is the tallest variety, has relatively few stolons, and is generally short-lived. Intermediate varieties have smaller leaves with greater stoloniferous growth, and are most resistant to stress. Low-growing clover ('weedy' or 'dutch' clover) has even smaller leaves and is largely not cultivated. We term these the Large, Intermediate, and Small varieties, respectively, below.

Population genomics dataset

The worldwide population genomics dataset presented here is an extension of the GLobal Urban Evolution Project (GLUE; *34*) with samples added from a key area for white clover domestication (Spain) as well as several cultivars. Full sampling methods for the GLUE project are included in (6), and we analyzed the subset of this original sampling that received low-coverage whole-genome sequencing. Briefly, we included 2616 samples collected between 2016-2019 from 50 different cities spanning the native range in Eurasia (12 cities) as well as introductions to North America (11 cities), South America (10 cities), Japan (four cities), China (four cities), Oceania (eight cities), and Africa (one city). Collections for these cities were made along urban to rural transects; however, we treat all collection sites from a single city as a single population as there is relatively little genetic differentiation among collection sites within each transect (7). Details for each of these populations can be found in Data S1. Samples sizes for each population ranged from 5-120. This heterogeneity in sample size was intentional as we wanted to include a number of cities with high sampling for better estimates of site frequency spectra and population-genomic statistics (31 cities; Ave. = 80.74, Std. = 17.7 individuals; *56*). We then added additional cities with lower sampling that we deemed as important areas for understanding colonization history (19 cities; Ave. = 5.95, Std. = 0.23 individuals). Environmental data for each sampling location was extracted from BIOCLIM using the *raster* v3.6-26 package in R and averaged across each city.

To supplement our GLUE sampling, we added 32 samples collected as seeds from 4 cities in Spain (A Corona, Granada, Salamanca, San Sebastian). These samples were also included in the fitness GWAS dataset below and were originally collected by Simon Innes in 2018 (9). Since we have low sample sizes in each city and genetic clustering analysis suggests similarity among these populations, we treat all populations from Spain as a single population. BIOCLIM data for Spain was extracted for each Spanish city in R (10) and we averaged each BIOCLIM variable over the four cities. Since there is likely substantial introgression from cultivar sequences in both native and introduced populations, we included 12 popular cultivars bred in the U.S. (Durana, Patriot, Renovation, Merit, Pilgrim, LA-S1, CA Ladino), Australia (Irrigation), and New Zealand (Crau, Grassland Huia, Grasslands Pitua). Note that some of these cultivars were derived from accessions collected far from where human-assisted breeding occurred (i.e. Crau was bred from lines collected in France and Pitua is a hybrid between Spanish and New Zealand collections; *29*). Cultivars include both Large and Intermediate varieties.

<u>Library construction and sequencing</u>

DNA extraction and library construction for GLUE samples has been previously described (6). However, molecular methods differed slightly for samples from Spain and cultivars. Briefly, we collected 1-3 trifoliate leaves per plant and either stored tissue at -80°C until extraction at University of Louisiana, Lafayette (ULL) or lyophilized tissue prior to shipping to ULL. DNA was extracted with a modified cetyltrimethylammonium bromide (CTAB) procedure using increased concentrations of CTAB and 2-β-mercaptoethanol (12, 13). DNA concentrations were quantified and diluted to 3ng/uL before undergoing a modified Nextera dual-index library preparation (14). We used Nextera XT Index Kit (Illumina; San Diego, CA USA) for barcoding libraries. Each library was quantified and concentrations normalized before pooling of 48 samples with unique barcodes into a single tube. Cultivars and samples from Spain were sequenced with the rest of the samples from the experimental common gardens described below (i.e. the Fitness GWAS). Each pool was sequenced by Novogene (Davis, CA) on a separate sequence lane using paired-end 150bp reads on a HiSeqX platform (14 total lanes). This resulted in low coverage across samples that was comparable to the GLUE genomic dataset (~1X). Notably, the Toronto population had been sequenced to ~10X coverage for the GLUE dataset and this difference in coverage from other samples can result in artifacts in population structure analysis (8). Thus, we downsampled the Toronto dataset using Samtools v1.10 (15) to ~2.5X coverage.

<u>Sequence processing, alignment, and filtering</u>

Sequence processing was identical for all samples included in these datasets (https://github.com/James-S-Santangelo/glue_dnaSeqQC). For each sample, we performed quality-control and examined sequences using FastQC v0.11.9 both on raw reads and on trimmed reads. We trimmed adapter sequencing and poly-g tails using the *-trim_poly_g* argument in *fastp* v023.2 (16). We aligned both reads from each individual to a haploid chromosome-scale reference genome (17) using *bwa mem* v0.7.17(18) with default settings. For each .sam file, we marked duplicated reads, coordinate-sorted, and converted to bam format using samtools v1.10(15). QC on each bamfile was performed using Qualimap v2.2.2 (19), Bamtools v2.5.1 (20), and multiQC v1.14 (21). From the reference genome and annotation, we constructed a list of four-fold degenerate sites using the Degeneracy Pipeline (https://github.com/tvkent/Degeneracy) to use for demographic analysis and converted to ANGSD format using an *awk* command (http://popgen.dk/angsd/index.php/Sites). In all of our analysis and text, we refer to chromosomes by number (1-16). This notation differs slightly from the reference genome which labels chromosomes by progenitor species (e.g. Chr01_Occ, Chr01_Pal). These terminologies can be easily linked (e.g. Chromosome 1 = Chr01_Occ, Chromosome 2 = Chr01_Pal, Chromosome 3 = Chr02_Occ, etc.).

<u>Analysis of demography and worldwide population structure</u>

We assessed population genomic diversity, differentiation, and structure using genotype likelihood approaches implemented in ANGSD v0.929 (22). To examine genetic diversity within each population, we first calculated genotype likelihoods and site allele frequency likelihoods (SAF) for each population independently using only 4-fold degenerate sites (-GL 1 -doMaf 2 -doCounts 1 -dumpCounts 2 -baq 2 -minQ 20 -minMapQ 30 -doSaf 1 -sites 4fold.sites). We conducted separate analyses on each chromosome for each population to better parallelize our computation. We calculated the folded, one-dimensional site frequency spectrum for each population with *realSFS* using the '-fold 1' option with 2000 iterations to ensure convergence (-maxIter). Site frequency spectrums were used to calculate thetas ($\theta_w$ and $\theta_\pi$) for each chromosome in each population using 'realSFS saf2theta' and the resulting output was used to calculate diversity statistics using 'thetaStat do_stat'. Theta and Tajima's D statistics were averaged across chromosomes using the number of sites on each chromosome. Average number of SNPs per population for these analyses was 10,784,068 (Std. 865,692).

Introductions are often expected to be accompanied by substantial bottlenecks and lower genetic diversity resulting from introduction of a limited number of individuals. However, admixture between genetically diverged native populations within the introduced range can result in greater genetic diversity and a limited genetic signature of a bottleneck. We examined whether populations of white clover in introduced areas have lower genetic diversity than in the native range using a one-way analysis of variance test implemented in R using the *oneway.test()* function that explicitly accounts for unequal variances using var.equal=FALSE (23). We conducted separate tests with each theta value as the response variable as $\theta_\pi$ is sensitive to the frequency differences in the site frequency spectrum while $\theta_w$ is not. We also conducted separate models using either all introduced population pooled vs native populations as levels of a factor or using each presumed independent introduction as levels of a factor (i.e., Africa, China, Japan, North America, Oceania, and South America as different levels). To detect potential bottlenecks following introduction, we conducted identical one-way ANOVAs with Tajima's D as the response variable. Bottlenecks (followed by population expansion) often result in overrepresentation of low frequency variants within the population (i.e. a negative Tajima's D value). We subsequently evaluated how different sample sizes and numbers of variable sites impacted calculations of diversity statistics and Tajima's D using linear regressions (*lm()* function in R). There were significant effects of both sample size and number of sites on $\theta_w$ (sample size: $r^2 = 0.13$ , $p = 0.01$; number of sites: $r^2 = 0.3$ , $p = 2.9e-5$ ) and Tajima's D (sample size: $r^2 = 0.22$ , $p = 0.0005$; number of sites: $r^2 = 0.52$ , $p = 1.5e-9$) presumably driven by better estimation of low frequency variants with larger sample sizes. However, variation in sample size and number of sites exists both in the native and introduced ranges and including either metric as a covariate in a model does not change the qualitative result. Thus, we elect to report the simpler models in the main text.

To examine demographic changes associated with colonization and introduction within each population in a coalescent framework, we used EPOS (24) to estimate Ne through time. Mutation rate was set to $1.8\times10^{-8}$ and each run included 1000 bootstrap iterations. Raw outputs were converted to a plottable format with the *epos2plot* function. We specifically examine the most recent 1000 years to determine whether we observe greater bottlenecks in introduced vs. native populations.

We investigated patterns of genetic differentiation within and among populations across the native and introduced ranges by calculating pairwise Fst values on four-fold degenerate sites in ANGSD. We recalculated SAFs for each population using the reference genome to assign major and minor alleles (-GL 1 -doMaf 2 -minMaf 0.05 -doCounts 1 -dumpCounts 2 -baq 2 -minQ 20 -minMapQ 30 -doSaf 5 -doMajorMinor 4 -sites 4fold.sites). Using these SAFs, we calculated 2d SFS between each population pair using 'realSFS'. Resulting 2D SFS were used in combination with the SAFs to generate fst binary files using 'realSFS fst index' and then weighted and unweighted measures of Fst (25, 26) were extracted using 'realSFS fst stats'. We designated each population pair as a comparison within the native range, within an introduced range, between a native and introduced range, or between populations from different introduced ranges. With a limited number of introductions to each introduced range, we expect that differentiation should be lower within introduced ranges. If populations in the introduced range were sourced from the same populations from the native range (or sourced through bridgehead events), we expect lower differentiation between populations from different introduced ranges as well as similar patterns of differentiation between each native:introduced range comparison. With repeated introductions and sharing between regions across the globe, we would expect genetic differentiation to be independent of our native vs. introduced population framework and to observe similar differentiation between populations from different introduced regions as between native and introduced populations.

To examine patterns of isolation by distance and isolation by environment in native and introduced ranges, we used pairwise Fst values within the native, North American, and South American ranges to conduct Mantel Tests. These tests were not conducted for other introduced regions as there were under five populations (Africa, China, Japan) or geographic distance matrices would not make sense over oceans (Australia and New Zealand in Oceania). Linearized pairwise Fst was calculated as Fst/(1-Fst). Haversine geographic distances between populations were calculated using *distm*() function within the *geodist* library (https://cran.r-project.org/web/packages/geodist/index.html). We calculated climatic distance between populations by using the *dist()* function in the *vegan* library (27) to calculate Euclidean distances between population mean annual temperatures and temperature seasonality (BIO1 and BIO4, respectively, from WorldClim dataset). Mantel tests (comparisons of linearized Fst matrixes with geographic distance matrices) were implemented using the *mantel*() function within the *vegan* library (27). We specifically used a Pearson correlation with 999 permutations to test statistical significance

We examined worldwide population structure and individual ancestry using NGSadmix (28). Unlike the population estimate of diversity and differentiation above, all samples were included in genotype likelihoods estimation using the parameters: -GL 1 -doGlf 2 -doMajorMinor 4 -doMaf 2 -doCounts 1 -baq 2 -minQ 20 -minMapQ 30 -minMaf 0.05 -sites 4-fold.sites. The resulting genotype likelihoods (i.e., beagle file) was to estimate individual ancestries using NGSadmix for three initial replicates of K=1-8 using 10,000 iterations per replicate (-maxIter). This analysis included 533,655 sites. The initial runs showed a single odd run (i.e. a highly divergent logLikelihood that we have been unable to repeat) for K=2. Thus, we re-ran 5 additional replicates at K= 2-4. To determine the most likely number of clusters, we examined standard deviations in likelihoods at each K and used the method of Evanno et al. (29) to identify the most likely number of ancestral clusters and the uppermost level of population structure. We visualized spatial variation in ancestry for most likely K value (K=3) by calculating population average ancestry for each population and plotting these values as pie charts on maps.

To further dissect population structure and examine introduction history within each introduced region, we used a principal component analysis. Using the same genotype likelihoods as within the NSGadmix analysis, we used PCAngsd (30) to generate a variance-covariance matrix of allele frequencies (pcangsd.py) and extracted the eigenvectors (i.e. the principal components) of the covariance matrix using *eigen()* function in R. Variance associated each principal component was assessed by dividing the eigenvalue for each PC axes by the sum of all eigenvalues. We calculated averages for PC1-5 for each population. We plotted population averages for native and introduced populations and identified the most closely related native populations to specific introduced populations as potential colonists. To examine potential clustering within the PCA by range (native/introduced or native/introduced/cultivar), we conducted PERMANOVA using the *adonis2*() function within the *vegan* library (27). We ran two separate models. First, to identify whether native and introduced populations exhibited clustering and differentiation, we used the first four PC axes as a response variable and used range (with native or pooled introduced populations as levels) as a factor. We also wanted to determine whether cultivars were differentiated from native and introduced populations. For this analysis, we ran a second identical PERMANOVA that included cultivars as well as native and introduced populations. Modeling was robust to the number of genetic PCs included. To test whether genetic PCs may reflect sampling limitations, we examined the association between each of the first four PCs and either number of individuals per population or coverage (*lm()* function). To determine whether PCs reflect isolation by environment, we examined associations between the first four PCs and several different BIOCLIM factors (e.g. mean annual temperature (BIO1), temperature seasonality (BIO4), mean annual precipitation (BIO12), precipitation in the coldest quarter (BIO19)). These were chosen because they were minimally correlated and represented ecologically meaningful climatic factors for white clover.

<u>Genome scans for local adaptation</u>
We examined regions of the genome affected by natural selection through two genome scans. Both genome scans rely on calculation of allele frequencies from genotype likelihoods for each population. We identified all sites that were polymorphic with a minor allele frequency greater than 5% across samples using ANGSD (-GL 1 -doGlf 2 -doMajorMinor 4 -doMaf 2 -baq 2 -minQ 20 -minMapQ 30 -SNP_pval 1e-6 -minMaf 0.05) in each range (Europe, North America, South America, Oceania, China and Japan) for climate adaptation scans or pair of ranges for contrast scans. We then called population allele frequencies for these sites in each population individually using ANGSD with no minor allele frequency filter (-GL 1 -doGlf 2 -doMajorMinor 4 -doMaf 2 -doCounts 1 -baq 2 -minQ 20 -minMapQ 30 -minMaf 0).

The first genome scan examined regions of the genome that were strongly differentiated between the native range and each of the five introduced ranges. These sites may be associated with phenotypes that facilitate invasiveness. We used the BayPass contrast statistic (31) to summarize allele frequency differentiation at each site between European populations and populations from an invasive range. Enrichment of contrast outliers was calculated for non-overlapping 20 kbp windows using the weighted-Z analysis (WZA)(32) and outlier windows were defined as the 1% tail of the distribution of WZA window scores.

The second genome scan tested for climate-related adaptation in the introduced ranges. To test for genomic regions with greater differentiation than expected by chance within each native range while accounting for genome wide population structure, we used the BayPass core model. Specifically, population allele frequencies were combined within ranges, including only sites with frequencies called in all populations (Europe: 22.7M; North America: 23.2M; South America: 20.7M; Oceania: 20.8M; China: 18.2M; Japan: 14.7M). We generated population covariance omega matrices for each range in BayPass v2.2 (31, 33) using 10,000 sites sampled from outside annotated genes, and then ran the BayPass core model to generate XtX statistics for each site. Next, correlations between population allele frequencies in each range and six minimally correlated bioclimatic variables (BIO1, BIO2, BIO8, BIO12, BIO15 and BIO19 from WorldClim dataset (10)) were quantified using the absolute value of Kendall's Tau. In each range, we used the weighted-Z analysis (WZA) to identify non-overlapping 20 kbp windows that were enriched for outliers for the XtX statistic and correlations with each bioclimatic variable. Outlier windows for each statistic were defined as the 1% tail of the distribution of WZA window scores. Outlier windows that overlapped between genome scans were identified, and their enrichment relative to a hypergeometric distribution was tested in R.

<u>Haploblock identification</u>
We identified population-genomic signatures of haploblocks using local principal component analysis, modifying the method described by Li and Ralph (34) to utilize covariance matrices from PCAngsd v1.10 (30), which were calculated in 100kbp windows from beagle files generated in ANGSD v0.929(22) (-GL 2 -doMajorMinor 1 -doCounts 1 -doGLF 2 -SNP_pval 1e-6 -doMaf 2 -doGeno -1 -doPost 1 -minMapQ 30 -minQ 20 -trim 5 -minMaf 0.05 -minInd 665 -geno_minDepth 2 -setMinDepthInd 2 -uniqueOnly 1). Local population structure along each chromosome was analyzed on five MDS axes and outliers were identified from the 5% corners of each pair of MDS axes.

Candidate haploblocks were identified by manual examination of MDS plots for each chromosome. Across each candidate region local PCAs were performed with ANGSD and PCAngsd using the same parameters used for the 100kbp windows. Heterozygosity was also calculated for each sample in each candidate region using ANGSD (-dosaf 1 -minMapQ 30 -minQ 20 -trim 5 -GL 2) and realSFS v0.929 (35) (-fold 1). After filtering out samples with less than 0.4X coverage, putative inversions were identified by the presence of three clusters of samples along a single principal component axis, indicative of two homozygous and one heterozygous inversion genotype, as well as by the presence of elevated heterozygosity in the region for samples genotyped as heterozygotes. We defined elevated heterozygosity as when the standard error of the mean (SEM) for each homozygote class did not overlap the SEM for the heterozygote class. We validated haploblock genotypes by performing LD scans with ngsLD v1.2.0 (--min_maf 0.05 --max_kb_dist 0) on 5000 randomly sampled sites from each chromosome containing a haploblock. For each haploblock, LD scans were run on a set of samples homozygous for the more common haploblock allele, as well as a random set of samples of the same size. To identify GO terms enriched in haploblocks, the R/topGO package (36) was used with Fisher's exact test, the 'weight01' algorithm, and a $p$-value < 0.05 to assess significance.

Associations between haploblocks and fitness

We examined patterns of local adaptation and the genomics of adaptation using four common gardens located in the southern and northern region of the native range (Montpellier, France and Uppsala, Sweden respectfully) and the southern and northern regions within the North American introduced range (Lafayette, USA and Mississauga, Canada respectfully). Common gardens were conducted for two years at each site, 2020-21 in North American gardens and 2021-22 in European gardens. Seedlings from the same lines were planted in each garden. Seeds were collected from 46 white clover populations spanning a 27° latitudinal gradient in Europe and from 47 additional populations spanning a 21° latitudinal gradient in North America. Field-collected seeds were all grown for a single generation in the greenhouse to minimize maternal effects, and then outcrossed within-populations by hand-pollination to create five outbred lines per population. Seeds from outbred lines were germinated and established for four weeks in 8cm peat pots within a greenhouse for each garden before being randomized and planted directly into the natural soil of a cultivated lawn at each site. Each grid had 1m spacing between all plants and the ground was covered with landscape fabric. Twelve cm holes were cut into the fabric for each plant, leaving a 2cm ring of natural vegetation of natural vegetation around each plant. Plants were given supplemental water for two weeks for establishment.

Multiple components of plant fitness were measured throughout the growing seasons of each location. In this study, we report two of the most critical measures – survival to flowering and total seed mass. Briefly, survival and flowering were surveyed weekly for each plant throughout the two-year experiment (except in colder climates when plants were covered by snow and/or dormant in the winter). Fruits were also collected weekly once plants reached maturity. All fruits were ground with a thresher (Precision Machine Co., Lincoln, NE, USA) and seeds were weighed to obtain a total seed set mass for each plant. Survival to flowering is a binary variable that indicates if a plant was able to flower during the two-year experiment and represents both viability and ability to mate. Total seed mass reflects both viability and fecundity as plants that did not produce any seeds had no seed mass.

Between 1-3 trifoliate leaves were harvested for DNA extraction from each surviving individual after 1-2 months of growth. In total, 656 samples (138 - Lafayette, Louisiana, USA; 327 - Mississauga, Ontario, Canada; 111 - Montpellier, France; 79 - Uppsala, Sweden) were sequenced for this experiment with 586 samples with sufficient coverage for GWAS analyses. DNA extraction, low coverage whole genome sequencing library construction, sequencing, and processing of raw reads to bam files were conducted as described above. To determine whether allelic variation in haploblocks was associated with differential fitness between gardens, we used our lcWGS to genotype haploblocks. To estimate haploblock genotypes in GWAS population samples, local PCAs were performed on each previously-identified haploblock region using BAM files from 2,660 GLUE samples as well as the 586 GWAS samples as described above. The first two principal components of covariance matrices for each haploblock region were visualized and used to assign GWAS samples to genotype clusters.

To assess whether haploblocks were associated with adaptation following introduction, we took a three-prong approach. First, we validated our GLUE data, by examining allele frequencies of haploblocks from populations collected across the native and introduced range. If adaptation is indeed occurring across climatic gradients, we would expect the independent sampling schemes to produce similar clinal gradients across gradients in putative agents of selection. Pooling samples originating from the same populations across the four gardens, we examined associations between latitude and allele frequency across North America and Europe. Linear models were conducted for each haploblock using the R/lm() function with number of haploblock alleles (0,1,2) as a response variable and latitude as an independent variable. Note that these models provide qualitatively similar results to generalized linear models using a binomial family and a logit link. We then qualitatively compared whether clines in the native and introduced region in these gardens matched the GLUE dataset.

Second, we examined how haploblock variation impacts fitness across gardens. If haploblock alleles underlie rapid adaptation, we expect to see higher fitness associated with alternative haploblock alleles in different gardens. Again, a hypothesis of which haploblock alleles should be favored in each garden can be derived from GLUE population genomic data. We model each of our two fitness metrics (survival to flower and total seed mass) in separate univariate generalized linear models for each haploblock to decipher which element of fitness particular haploblocks impact. Fitness measures were response variables while Garden, Genotype, and Garden:Genotype interaction were included as factors. GLMs were implemented using *glm()* and statistical significance of each factor was assessed using *Anova()* with Type III sum of squares in the *car* library (37). Survival to flower was modeled within binomial error distribution and a logit link. Total seed mass was log(+1) transformed and modeled with gaussian distribution and identity link (i.e., a general linear model). We created detailed expectations for how selection should act in each population based on the haploblock allele frequencies in populations within North America and Europe calculated from our large population genomics dataset above. For instance, the alternative allele for HB13 is at high frequency in both Spanish and Swedish populations, but at low frequency in both high and low latitude North American populations, thus, we would expect that the alternative allele would have higher fitness in both European Gardens and the reference allele would have higher fitness in both North American gardens.

Finally, we also calculated relative fitness from total seed mass data for each haploblock to better understand the strength of selection acting on each haploblock within each garden. Relative fitness for each haploblock was calculated by dividing each individual value for total seed mass by the average value of total seed mass for the genotype with the highest fitness in the garden.

Association mapping within haploblocks

While associations between allelic variation in haploblock and fitness are indicative of adaptation, associations between individuals SNPs within haploblocks with fitness provide insight into the genes and phenotypes potentially under selection. We conducted association mapping with a genotype-likelihood framework implemented in ANGSD. First, we conducted independent GWAS for the two fitness traits in each garden. Second, we pooled European gardens and North American Gardens and conducted GWAS for each trait in each pooled sample. Genotype likelihoods were estimated for each range using 569 contemporary white clover samples (174 individuals from the native European range and 395 individuals from the introduced North American range) in ANGSD (-GL 1 -minMaf 0.05 -minMapQ 30 -minQ 20).

To perform each genome wide association, we employed a hybrid model (-doAsso 5) that first uses a score statistic to evaluate the joint maximum likelihood estimate between a trait and an observed marker

(38). If the chi-square test falls below a particular threshold (-hybridThres 0.05), a latent genotype model with an expectation-maximization (EM) algorithm was then performed that maximizes the trait-marker likelihood using a weighted least squares regression (39). With the EM algorithm, effect sizes ($\beta$) can be estimated and reported. To control for the effect of population structure within each range, a covariance matrix of estimated allele frequencies was constructed using *PCAngsd* (30). The top 20 principal components were calculated from the matrix and then added to the GWAS model as covariates. In the North American and European combined GWAS analyses, we added garden as a covariate to control for the effect of different growing environments at either common garden. We included a flag to include only SNPs that we were confident were polymorphic (-SNP_pval 1e-6). We corrected the alpha value using a conservative Bonferroni correction using the number of SNPs at a particular haploblock as the assumed number of multiple comparisons. To examine whether the number of fitness GWAS hits exceeded the genome-wide average, we conducted permutation analyses where the expected number of hits were simulated by taking random sequence lengths across the genome equal to the base pair length of each haploblock 10,000 times per chromosome (160,000 total samples) with replacement and then extracting the number of the number of GWAS hits for each sample. P-values were obtained as the ratio of number of sampled regions with more hits than within the target haploblock divided by total number of sampled regions. The significance threshold for categorizing a SNP as a hit within the randomly sampled sequences was equal to the Bonferroni correction applied to each haploblock region of a particular trait and garden.

Differential Expression Analysis

We performed a manipulative experiment to examine variation in expression between white clover populations in native and introduced range under dry-down and well-watered conditions. We selected 1-3 maternal lines from each of 3-4 populations from low latitude and high latitude in the European and North American ranges (14 total populations; 47 total samples).

We scarified seeds from each line and planted them in 4' pots filled with PRO-MIX LP-15 soil pre-saturated with water. Pots were placed in AR-66L2 growth chambers (Percival Scientific; Perry, IA USA) and misted daily to promote germination. Growth chambers were set to a constant 22°C with a 14hr:10hr day:night cycle. Germinants were transplanted to new plants randomized within flats that would eventually become control and dry down treatments. Flats were rotated within chambers every three days to minimize microenvironmental variation. Plants were grown for six weeks to accumulate above and below ground biomass. At six weeks, all pots were saturated with water by bottom-watering. Number of leaves on each plant were counted and one plant was excluded from further analysis because it was substantially larger than the other plants. Plants in the control (well-watered) flats received periodic watering according to our standard greenhouse protocol. Plants in the dry down treatment did not receive additional water. Each day, we assessed soil moisture in each pot using a SMT150T soil moisture meter (Dynamax; Houston TX). Control and dry down treatments functioned as expected with separation in volumetric water content between treatments after control plants were watered on day 9.

Leaf tissue from two healthy adult leaves was flash frozen in liquid nitrogen ten days after the dry down treatment began from plants in both the well-watered and control treatment. Total RNA was extracted with Direct-zol RNA Miniprep (Zymo; Irvine, CA USA). RNAseq libraries were constructed with QuantSeq 3' mRNA-Seq FWD prep kit (Lexogen; Vienna, Austria) using an input of ~415ng total RNA/sample following manufacturers protocols. Two samples had low RNA concentrations and we followed Lexogen's modified procedure for low input samples. All individuals were barcoded and multiplexed in a single tube. We sequenced this library on a single HiSeqX lane (PE 150bp reads) through Novogene (Sacramento, CA USA). A second round of multiplexing and sequencing was conducted for a subset of samples that had low data yield the first round, and files from both rounds were subsequently merged. Individuals averaged 37,241,613 reads (sd: 16,534,847) with 444,196 reads on average effectively mapping (sd: 243,429) due to DNA contamination.

We used *fastp* v023.4 (16) to trim adapters and poly-a tails. Microbial RNA contamination was removed by aligning each sample to all fungal and bacterial assembly genomes within the NCBI database using *bowtie2* v2.5.1 (40). The bacterial and fungal genome database was indexed with the 2.2.4 release of *bowtie2*. The remaining reads that did not align to the contamination database were presumed to be white clover RNA reads. A transcriptome was created from the *T. repens* reference genome using *gffread* v0.12.7 (41). A full decoy-aware transcriptome was constructed to mitigate specious mapping of reads that occur from unannotated genomic loci that are similar to the annotated transcriptome. A mapping-based index was then constructed using a k-mer hash over k-mers of length 31 using *Salmon* v1.10.2 (42). Read mapping to the transcriptome and quantification was then performed with *Salmon* v1.10.2[41] in mapping-based mode using the *T. repens* transcriptome and RNA sequences. To test for differences in transcript abundance between dry down and well-watered treatment groups, between the North American and European range, and between high and low latitude populations, we used *DESeq2* (43). We used two different models to examine differential patterns of gene expression across treatments, range and latitude. The first included all interactions (treatment*range*latitude). The second set of models were univariate models examining differential expression across treatment, range and latitude separately. False discovery rate (FDR) for each gene was calculated and a gene was then categorized as differentially expressed if FDR was < 0.1.

We evaluated whether transcribed genes located within haploblocks were more or less differentially expressed than in other regions of the genome through a resampling analysis. The mean absolute log2FoldChange of all genes within each haplotype was compared to a simulated null distribution. The null distribution was constructed by randomly sampling 10,000 different syntenic genomic regions containing the number of genes equal to the number found in each haploblock with replacement, and then calculating the mean absolute log2FoldChange for each genomic region. The probability of sampling the observed expression level of each haploblock from the null distribution was evaluated as the ratio of the number of simulated regions exceeding the observed haploblock mean absolute log2FoldChange divided by total number of simulations. This process was repeated for each comparison (range, treatment, latitude).

**Supplemental Materials and Methods References**

1. A. G. Griffiths, *et al.*, Breaking free: The genomics of allopolyploidy-facilitated niche expansion in white clover. *Plant Cell* **31**, 1466–1487 (2019).

2. T. Kjærgaard, A plant that changed the world: The rise and fall of clover 1000-2000. *Landscape Research* **28**, 41–49 (2003).

3. L. Carrier, K. S. Bort, The history of Kentucky bluegrass and white clover in the United States. *Agronomy Journal* **8**, 256–267 (1916).

4. J. R. Caradus, A. C. MacKay, D. R. Woodfield, J. Van Den Bosch, S. Wewala, Classification of a world collection of white clover cultivars. *Euphytica* **42**, 183–196 (1989).

5. D. Ogle, L. St. John, Ogle, D., St. John, L. 2008. Plant Guide for white clover (Trifolium repens L.). ,. (2008).

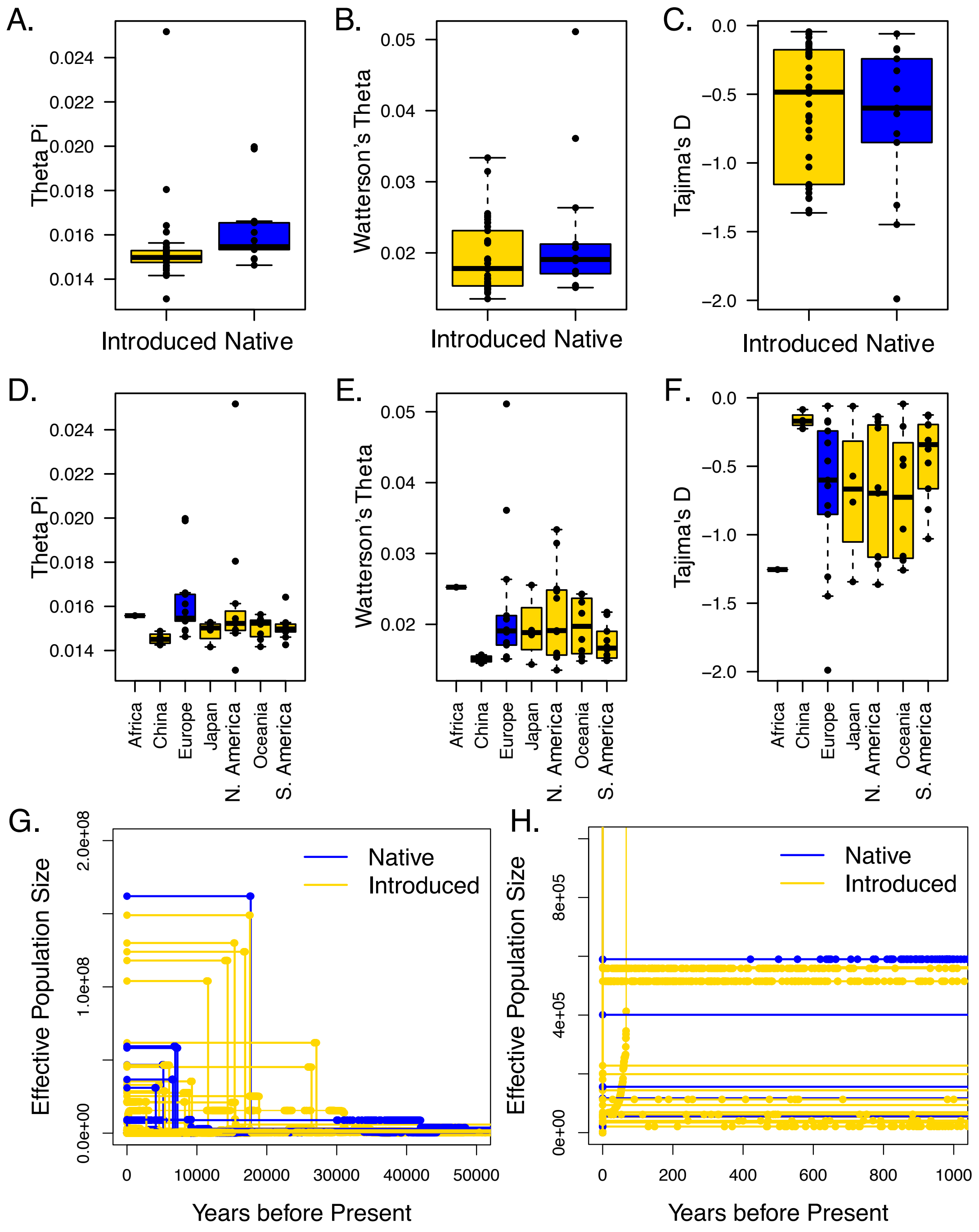

**Fig. S1. Population genetic summary statistics across native and introduced ranges. (A-C)** Boxplots compare measures of genetic diversity ($\pi$ and $\theta_w$) as well as Tajima's D between native and introduced ranges. **(D-F)** Boxplots further parse introduced population into individual introduction events. Each point represents the genome-wide average for a single population. Box edges in boxplots represent the interquartile range, center line represents the median, and upper and lower whiskers are the largest value either greater or less than, respectively, 1.5 times the interquartile range. **(G,H)** Coalescent reconstructions of effective population size through time as estimated through EPOS[52]. Neither native nor introduced populations exhibit any signatures of bottlenecks following introduction. Instead, most populations show signs of population expansion.

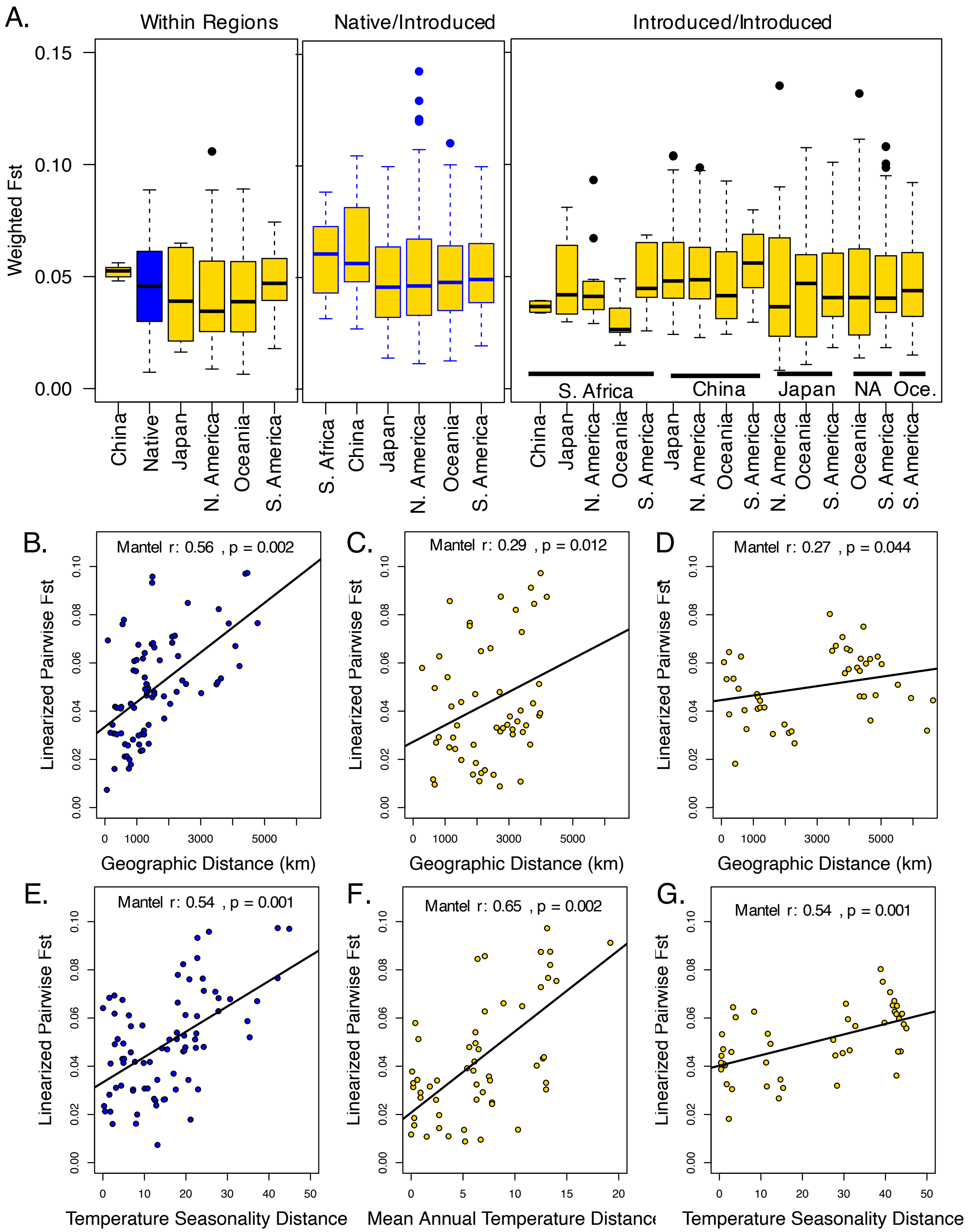

**Fig. S2. Population structure within and among native and introduced ranges. A.** Weighted pairwise fst within each range, weighted pairwise fst values between native and introduced populations, and weighted pairwise fst values between populations in different introduced ranges. Pairwise fst values are generally low and similar across all worldwide populations. **B-G.** Mantel tests for isolation by distance (B-D) and isolation by environment (E-G) across the native range (B,E), North America (C,F) and South America (D,G).

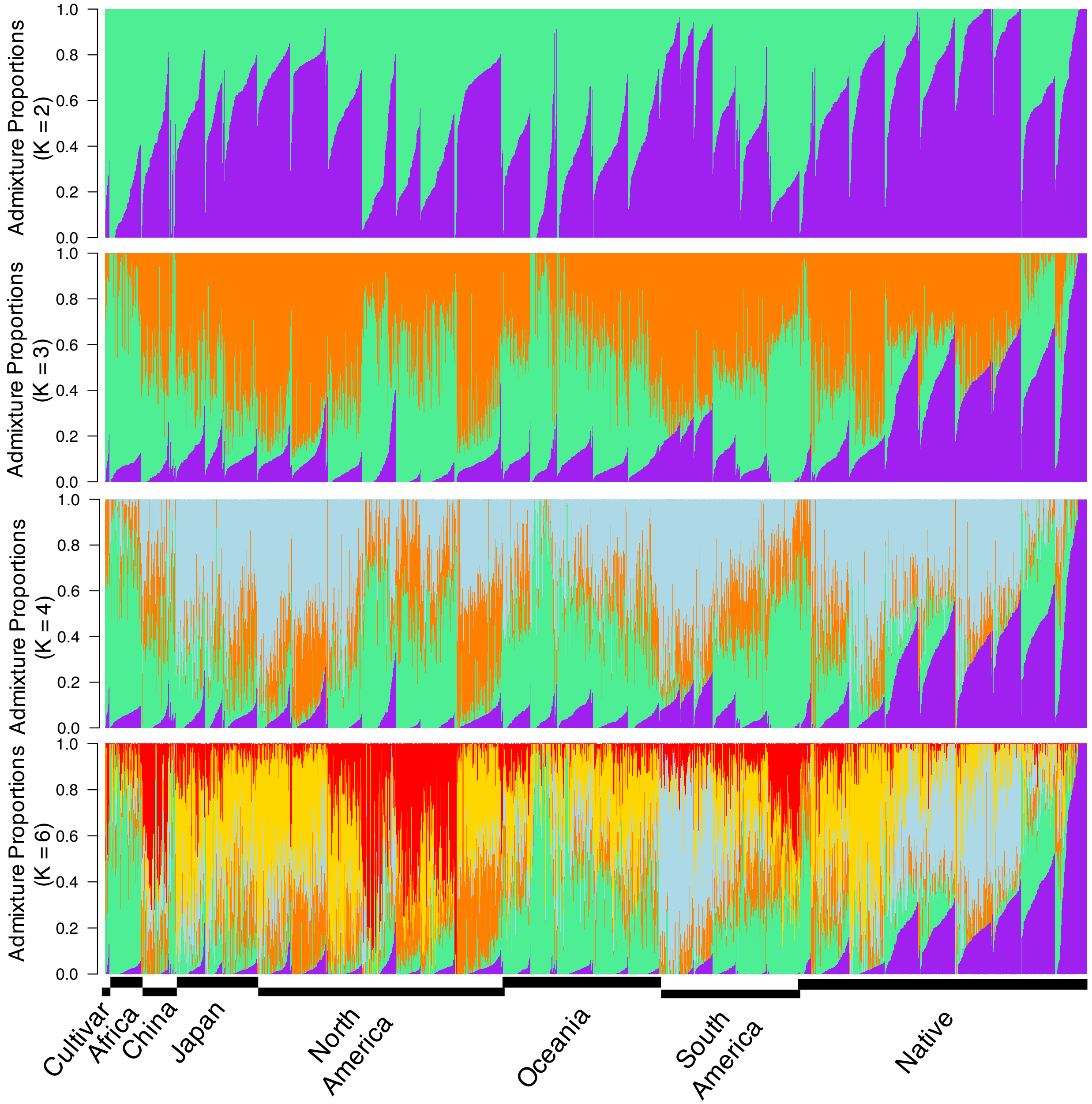

**Fig. S3. NGSadmix assumed population ancestry mapped across worldwide sampling**. Barplots depicts ancestry output from K=2,3,4, and 6 K-values. Best K was K=3. Individuals are organized along the x-axis by population sorted by continent, longitude, and ancestry values.

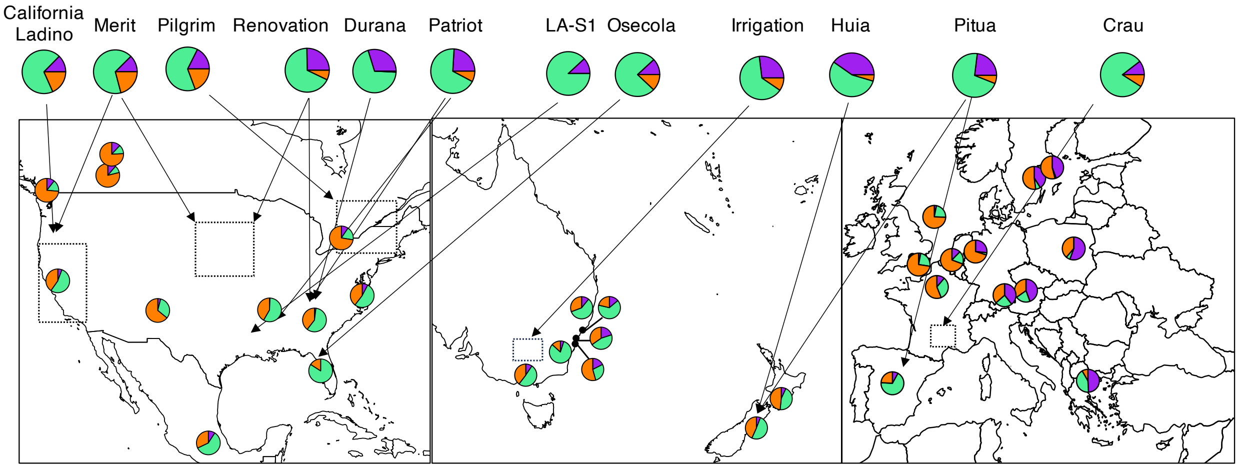

**Fig. S4. Cultivar assumed population ancestries relative to wild population sampling**. Each pie chart within map inserts reflects the average ancestries (K=3) from a given population. Dotted lines reflect relative locations where the wild stock was collected to generate each cultivar.

A.

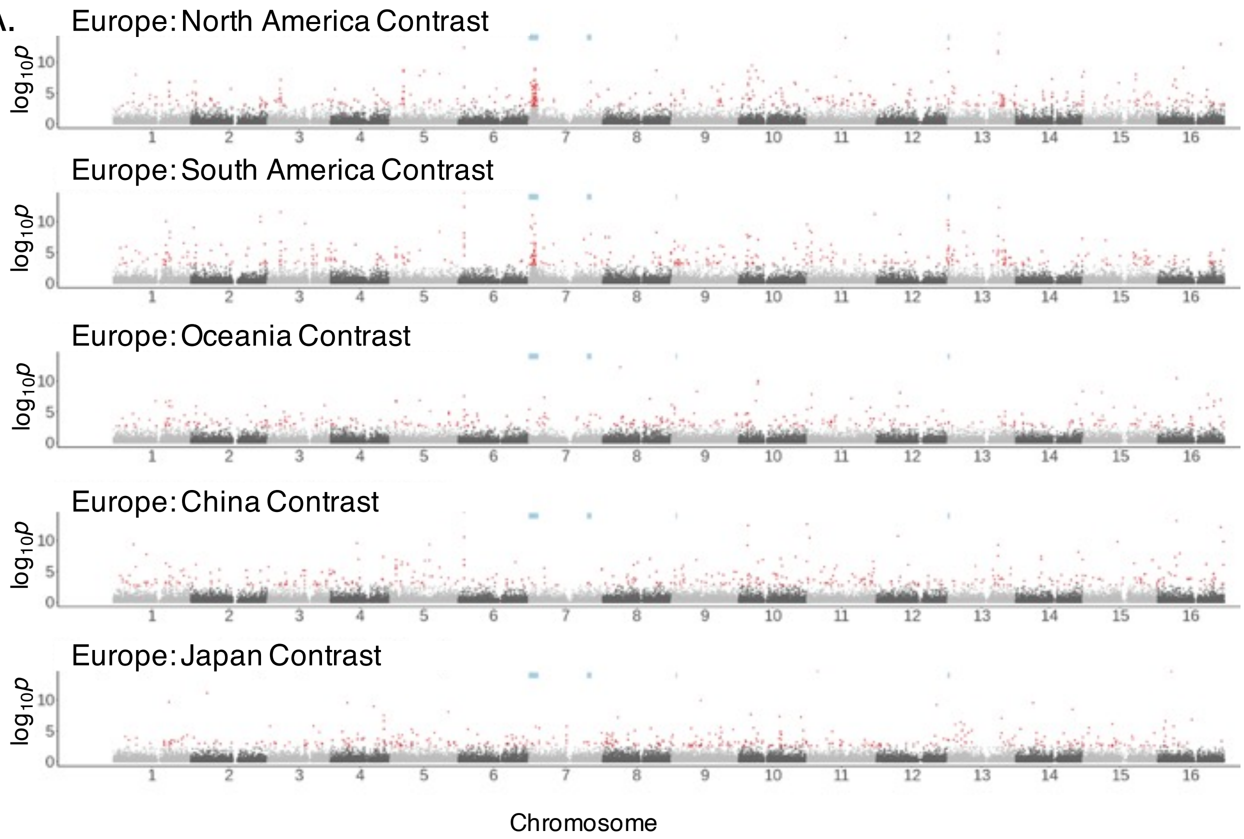

B.

| Native: Invasive Comparison | contrast | EU | | Invasive Region | |
|---|---|---|---|---|---|
| | | XtX | XtX-EAA | XtX | XtX-EAA |
| Europe:North America | 443 | 50 | 29 | 48 | 26 |
| Europe:South America | 443 | 56 | 35 | 48 | 23 |
| Europe: Oceania | 397 | 33 | 19 | 22 | 6 |
| Europe: China | 441 | 43 | 28 | 30 | 9 |
| Europe:Japan | 443 | 4 | 2 | 5 | 1 |

**Fig. S5. Genome scan for differentiated regions between Europe and each invasive range**. **A.** WZA 20 kbp window *p*-values against genomic location for contrast between Europe and each invasive range. Red points indicate the 1% tail of WZA scores. Blue bars indicate haploblock locations. **B.** Overlap between outlier 20kbp windows significant within the contrast analyses and outlier 20kbp windows within each introduced range (XtX) or across climatic gradients within each range (XtX-EAA).

A.

| All | Bio1 | Bio2 | Bio8 | Bio12 | Bio15 | Bio19 |
|---|---|---|---|---|---|---|
| HB7a1 | 0.139 | 0.013 | 0.263 | 0.188 | 0.022 | 0.045 |
| HB7a2 | 0.009 | 0.259 | 0.173 | 0.149 | 0.336 | -0.06 |
| HB7b | -0.17 | 0.024 | -0.2 | 0.007 | -0.04 | 0.096 |
| HB9 | -0.05 | 0.022 | 0.011 | -0.07 | 0.008 | -0.09 |
| HB13 | 0.099 | 0.032 | 0.113 | 0.101 | 0.102 | -0.01 |

| Europe | Bio1 | Bio2 | Bio8 | Bio12 | Bio15 | Bio19 |
|---|---|---|---|---|---|---|
| HB7a1 | -0.5 | 0.046 | 0.595 | -0.14 | 0.351 | -0.63 |
| HB7a2 | -0.08 | 0.295 | 0.357 | 0.388 | 0.171 | 0.016 |
| HB7b | 0.107 | -0.17 | -0.23 | 0.443 | -0.11 | 0.534 |
| HB9 | 0.242 | 0.545 | 0.061 | 0.121 | 0.182 | 0.061 |
| HB13 | -0.44 | -0.11 | 0.137 | -0.23 | 0.321 | -0.26 |

| North America | Bio1 | Bio2 | Bio8 | Bio12 | Bio15 | Bio19 |
|---|---|---|---|---|---|---|
| HB7a1 | 0.273 | 0.236 | 0.273 | -0.16 | 0.164 | -0.13 |
| HB7a2 | -0.24 | 0.018 | -0.02 | -0.09 | 0.236 | -0.35 |
| HB7b | -0.29 | 0.212 | -0.14 | -0.25 | -0.14 | -0.21 |
| HB9 | 0.367 | 0.257 | -0.15 | 0.073 | -0.15 | -0.04 |
| HB13 | 0.055 | 0.164 | -0.24 | -0.02 | 0.164 | 0.091 |

| South America | Bio1 | Bio2 | Bio8 | Bio12 | Bio15 | Bio19 |
|---|---|---|---|---|---|---|
| HB7a1 | 0.227 | -0.13 | 0.378 | 0.428 | -0.03 | 0.025 |
| HB7a2 | 0.244 | -0.2 | 0.556 | 0.333 | -0.11 | -0.29 |
| HB7b | -0.2 | 0.422 | -0.42 | -0.2 | 0.333 | 0.333 |
| HB9 | -0.18 | 0.09 | -0.23 | -0.05 | 0.09 | 0.045 |
| HB13 | 0 | -0.36 | 0.225 | 0.315 | -0.27 | -0.14 |

| Oceania | Bio1 | Bio2 | Bio8 | Bio12 | Bio15 | Bio19 |
|---|---|---|---|---|---|---|
| HB7a1 | 0.571 | 0.429 | 0.857 | 0.214 | 0.571 | 0.071 |
| HB7a2 | -0.18 | -0.11 | -0.04 | -0.11 | 0.182 | 0.036 |
| HB7b | -0.57 | -0.04 | -0.42 | -0.79 | -0.19 | -0.49 |
| HB9 | -0.19 | 0.189 | -0.04 | 0.113 | -0.11 | -0.19 |
| HB13 | 0.5 | 0.357 | 0.786 | 0.429 | 0.5 | 0.143 |

| China | Bio1 | Bio2 | Bio8 | Bio12 | Bio15 | Bio19 |
|---|---|---|---|---|---|---|
| HB7a1 | 0.236 | -0.24 | -0.24 | 0.236 | 0.236 | 0.236 |
| HB7a2 | -0.18 | 0.183 | -0.55 | -0.18 | 0.183 | -0.18 |
| HB7b | 0.236 | -0.24 | -0.24 | 0.236 | 0.236 | 0.236 |
| HB9 | 0.183 | -0.18 | 0.548 | 0.183 | -0.55 | 0.183 |
| HB13 | -1 | 1 | -0.67 | -1 | 0.333 | -1 |

| Japan | Bio1 | Bio2 | Bio8 | Bio12 | Bio15 | Bio19 |
|---|---|---|---|---|---|---|
| HB7a1 | -0.33 | -0.67 | -0.33 | 0 | 0 | 0.333 |
| HB7a2 | 0 | 0.333 | 0 | -0.33 | -0.33 | 0 |
| HB7b | 0 | -0.33 | 0 | 0.333 | -0.33 | 0.667 |
| HB9 | -0.67 | -1 | -0.67 | -0.33 | -0.33 | 0.667 |
| HB13 | -0.67 | -0.33 | -0.67 | -1 | -0.33 | 0 |

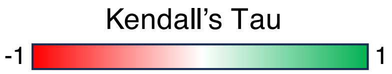

B.

| Region | XtX | Bio1 | Bio2 | Bio8 | Bio12 | Bio15 | Bio19 | All |
|---|---|---|---|---|---|---|---|---|
| Europe | 446 | 82 | 75 | 66 | 32 | 52 | 72 | 232 |
| North America | 449 | 108 | 18 | 41 | 48 | 17 | 29 | 207 |
| South America | 448 | 27 | 30 | 56 | 71 | 10 | 17 | 140 |
| Oceania | 400 | 21 | 55 | 27 | 22 | 27 | 19 | 116 |
| China | 448 | 26 | 26 | 32 | 26 | 16 | 26 | 69 |
| Japan | 449 | 28 | 30 | 28 | 27 | 32 | 23 | 110 |

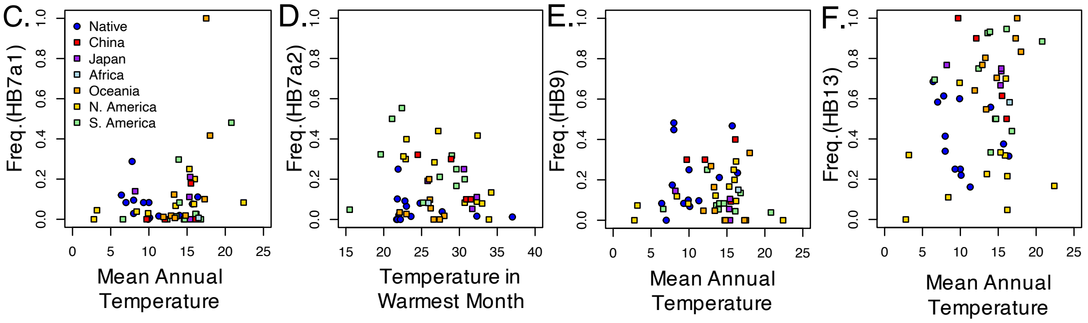

**Fig. S6. Climatic correlations with haploblocks and genome-wide variation. A**. Associations between haploblocks and climatic variables from the WORLDCLIM database. **B.** Overlap between outlier 20kbp XtX windows within each introduced range and outlier windows associated with each climatic variable. Abbreviations: Bio1: Annual Mean Temperature, Bio2: Mean Diurnal Range, Bio8: Temperature in the Wettest Quarter, Bio12: Annual Precipitation, Bio15: Precipitation Seasonality, Bio19: Precipitation in the Coldest Quarter.

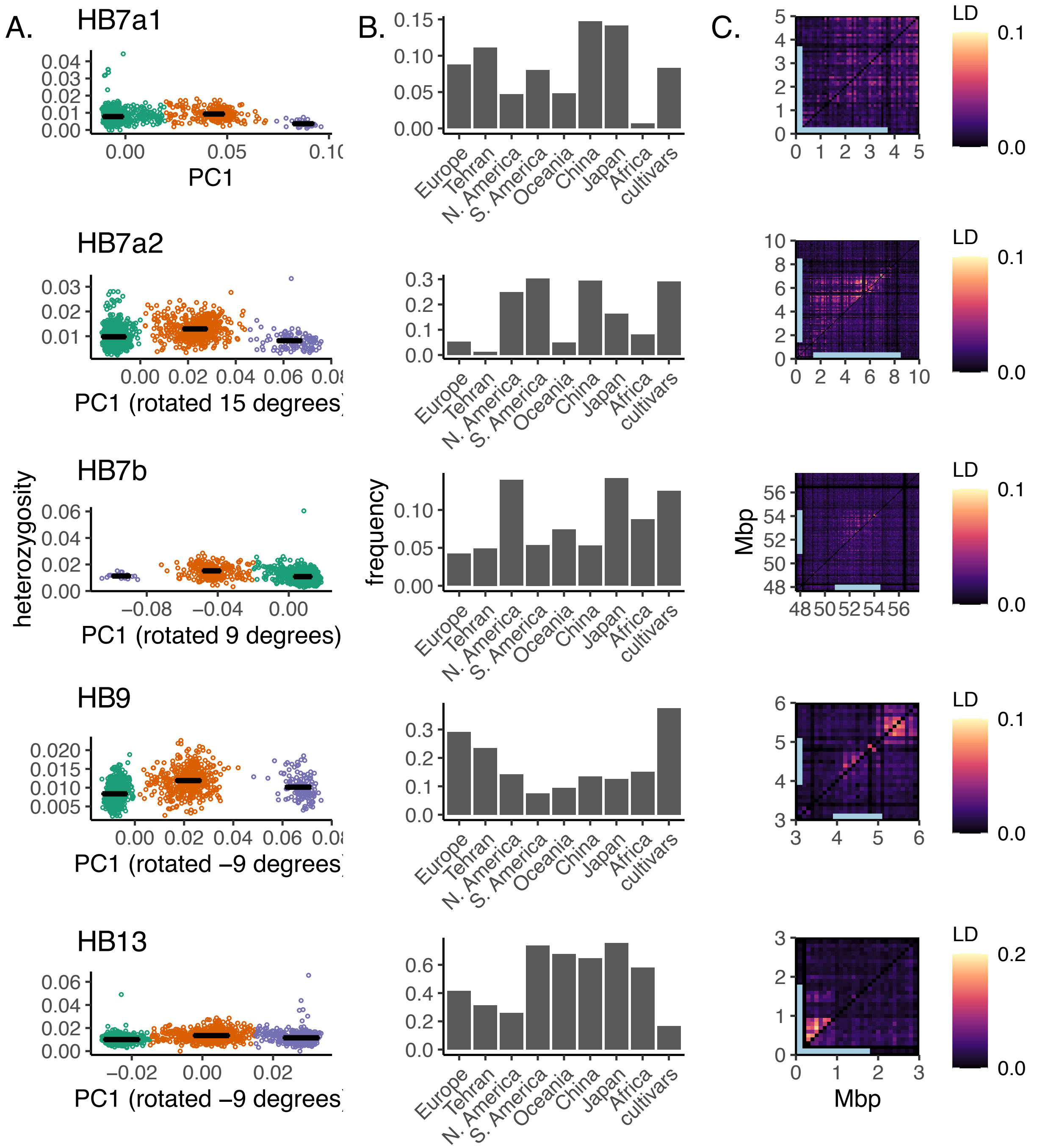

**Fig. S7. Five haploblocks–population-genomic signatures of structural variants**. **A.** Three clusters indicative of two homozygous (green and purple) and one heterozygous (orange) structural variant genotypes separate along the first principal component of genetic variation across each haploblock, and furthermore putative heterozygotes show significantly elevated heterozygosity (boxes denote mean +/- SEM for each cluster). **B.** Estimated allele frequencies of each haploblock. **C.** Local patterns of linkage disequilibrium (the second highest value in each 100kb window) corresponding to haploblock regions (blue bars) are present in a random sample of individuals (top triangle; matching sample size of bottom triangle) but absent in samples homozygous for the common haploblock allele (bottom triangle).

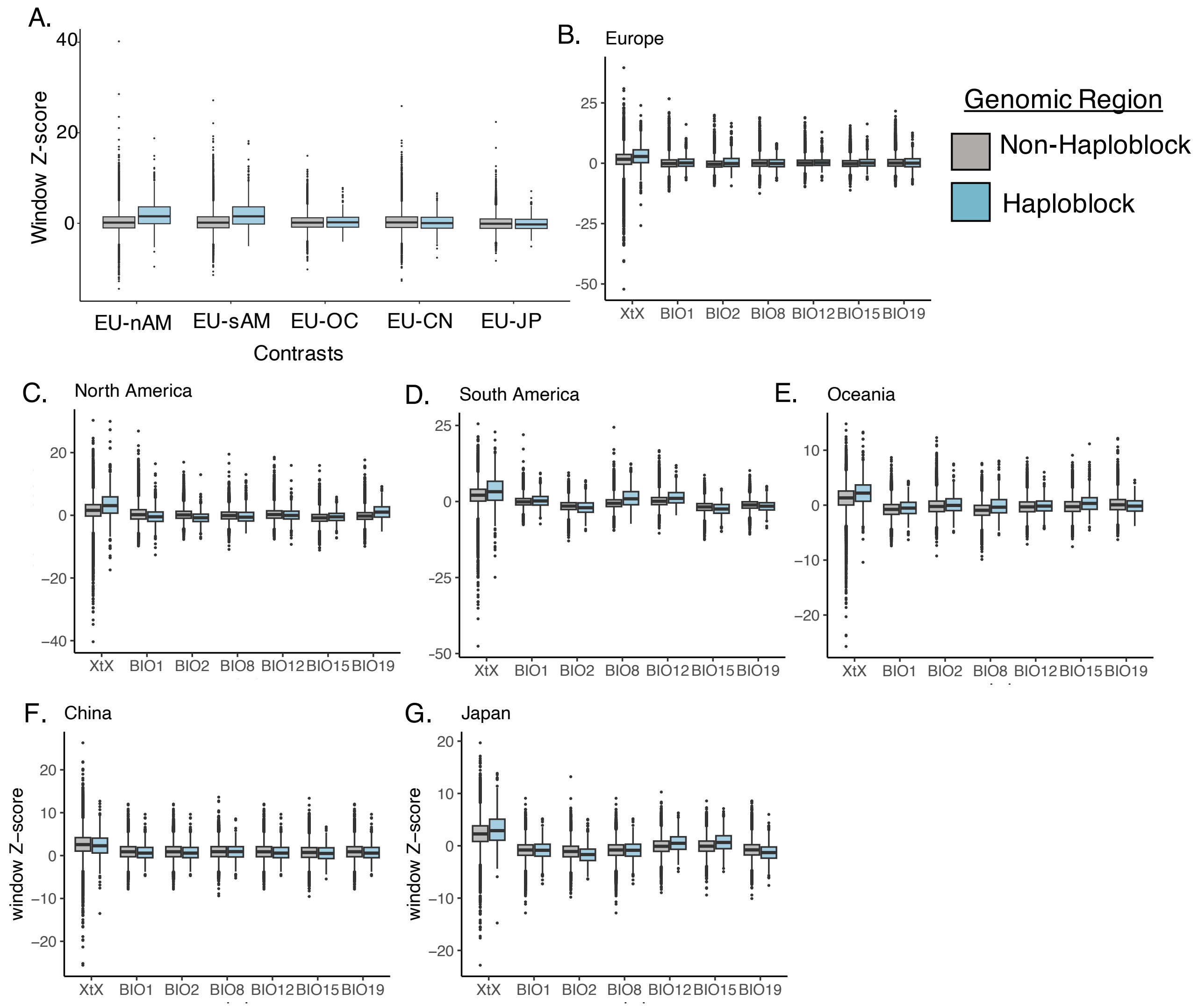

**Fig. S8. Comparisons of selective signatures within haploblock and non-haploblock windows across the genome. A.** Distribution of WZA 20 kbp window scores for contrasts between Europe and each invasive range for non-haploblock (gray) and haploblock (light blue) windows. **B-G.** Distribution of XtX statistics and Kendall's Tau for several climatic variables from the WorldClim dataset for each region. For boxplots, box edges represent the interquartile range, the center line in the box is the median, and the whiskers represent 1.5 times less or greater than the interquartile range. Abbreviations: Bio1: Annual Mean Temperature, Bio2: Mean Diurnal Range, Bio8: Temperature in the Wettest Quarter, Bio12: Annual Precipitation, Bio15: Precipitation Seasonality, Bio19: Precipitation in the Coldest Quarter. EU = Europe, nAM = North America, sAM = South America, OC = Oceania, CN = China, and JP = Japan.

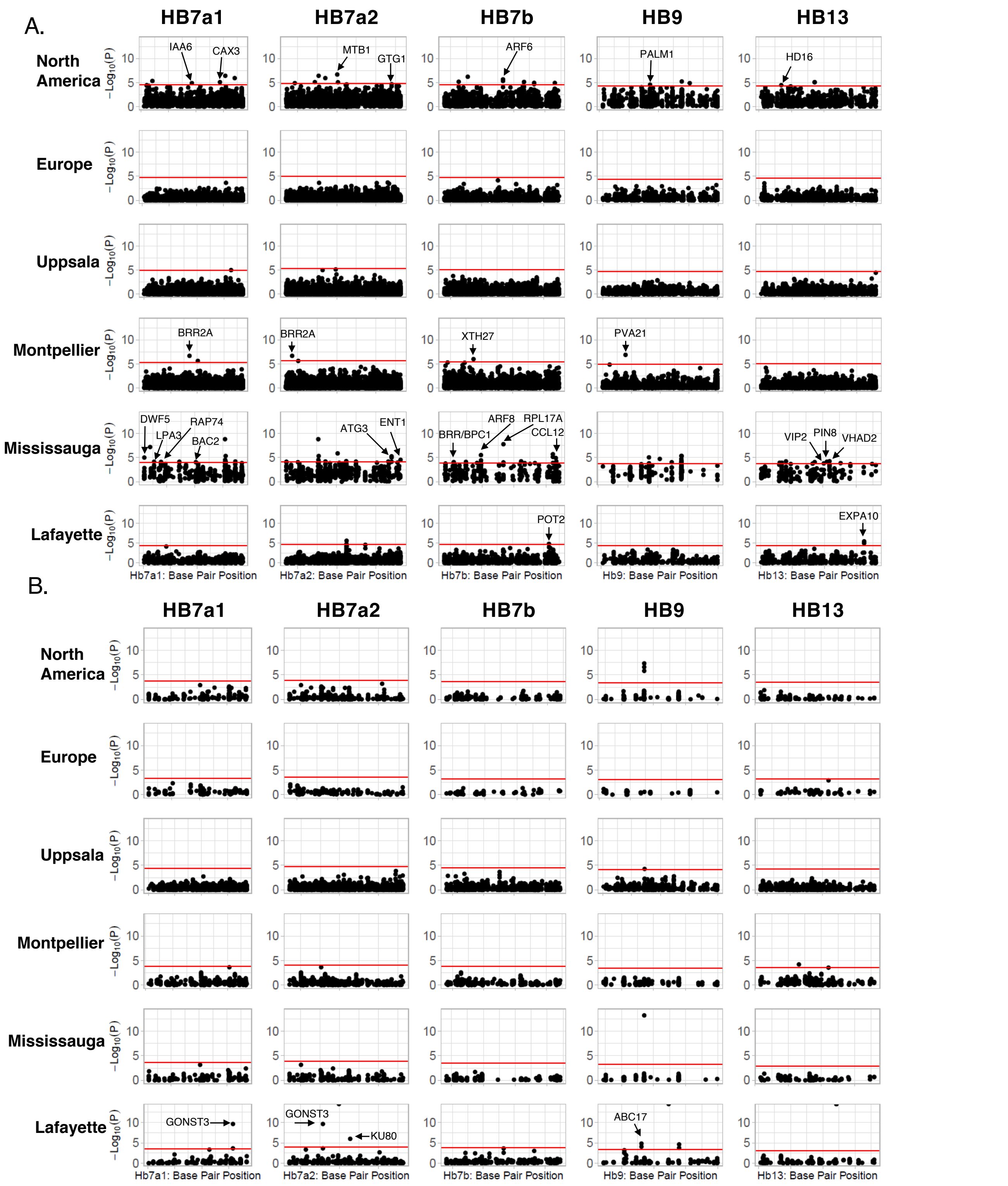

**Fig. S9 Manhattan plots summarizing associations with fitness across haploblocks**. **A.** Associations between haploblock genotypes with survival to flowering. **B.** Associations between haploblock genotypes with total seed mass (including zeros for individuals that did not survive to flowering). The Bonferroni corrected significance threshold (horizontal red line) is specific to each haploblock and garden. Gene names are given only for hits landing within coding sequence of annotated genes.

A.

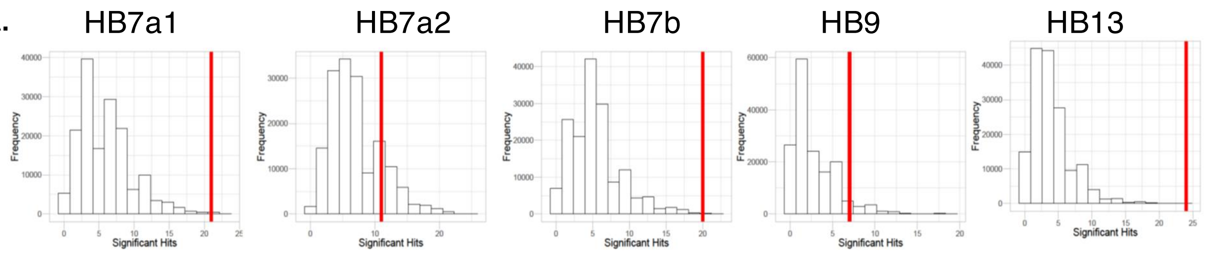

B.

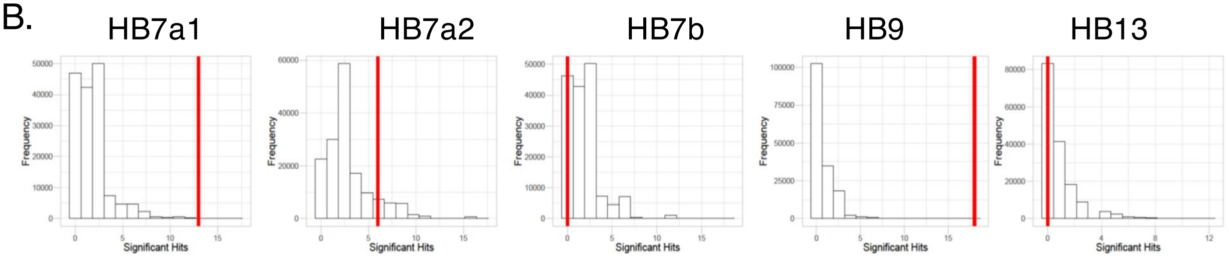

C.

| Region | Differentially Expressed Genes (%) | | | Absolute Expression (log2FoldChange) | | |
|---|---|---|---|---|---|---|
| | Range | Treatment | Latitude | Range | Treatment | Latitude |
| Genome-Wide | 0.205 | 0.129 | 0.215 | - | - | - |
| Hb7a1 | 0.207 | 0.114 | 0.205 | 6.06 / 7.06 | 8.1 / 8.9 | 6.08 / 7.26 |
| Hb7a2 | 0.394 | 0.217 | 0.404 | 6.82 / 7.06 | 8.43 / 8.9 | 6.88 / 7.26 |
| Hb7b | 0.142 | 0.081 | 0.136 | 7.14 / 7.06 | 8.28 / 8.88 | 6.53 / 7.26 |
| Hb9 | 0.053 | 0.035 | 0.053 | 5.44 / 7.04 | 7.58 / 8.88 | 5.71 / 7.27 |
| Hb13 | 0.087 | 0.043 | 0.093 | 7.48 / 7.07 | 8.94 / 8.87 | 7.8 / 7.26 |

**Fig. S10**

**Characterization of the genic content of haploblock regions**. **A-B.** Histograms summarizing expected numbers of GWAS hits for each haploblock for survival to flowering (A) and total seed mass (B) within the North American common gardens across 160,000 simulations. Observed number of GWAS hits are displayed as vertical red lines. **C.** Differential expression analysis of RNAseq data within each haploblock. Percentage of genes differentially expressed and absolute expression ($log_2$FoldChange) is presented between ranges (Europe vs. North America), between Treatments (Well-Watered vs. Dry Down), and between Latitudes (Low vs. High). Values on either side of paratheses for absolute expression are the observed / expected values. Expected values are derived from a permutation analysis that re-sampled regions from across the genome with replacement. P-values are greater than 0.05 in all cases.

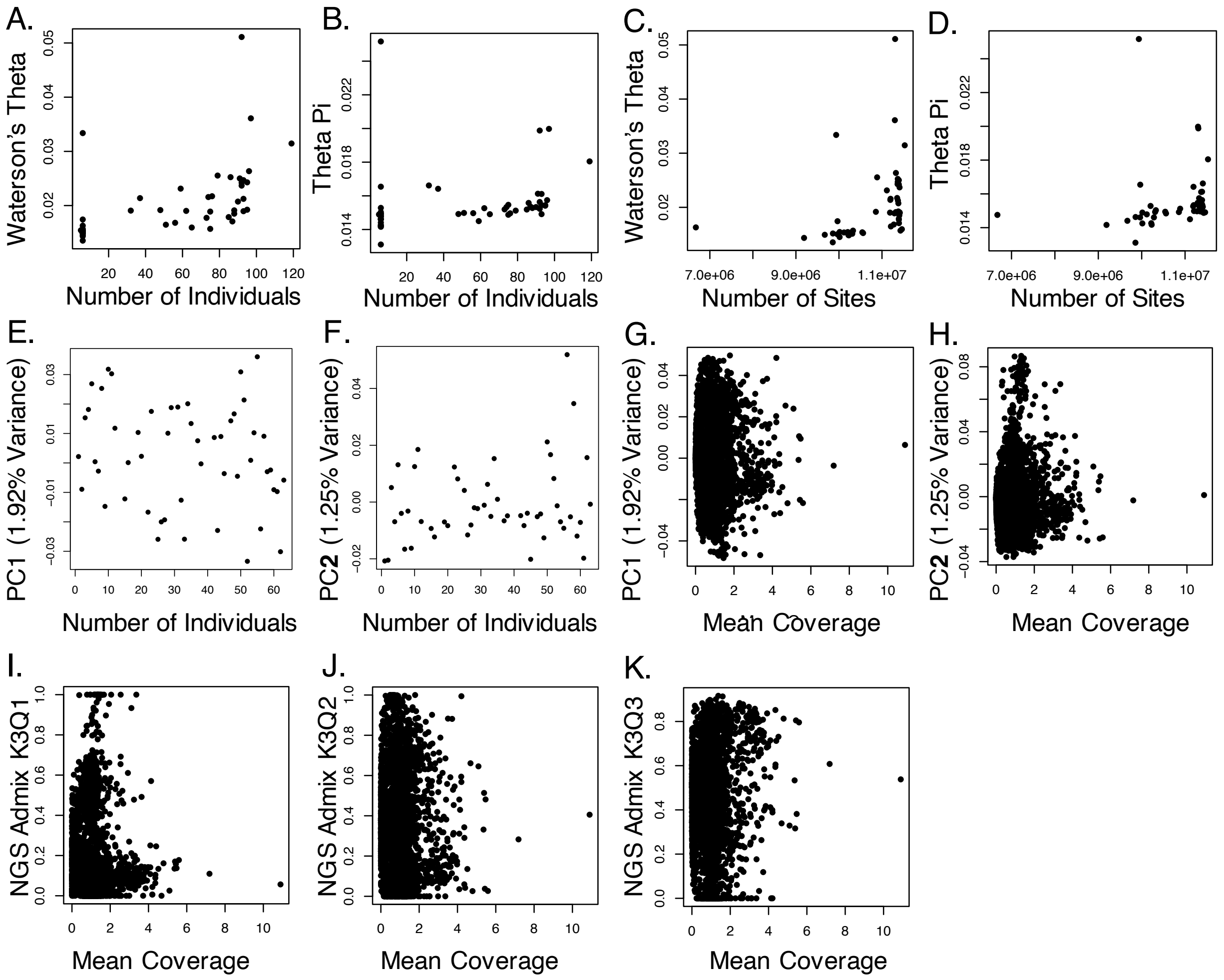

**Fig. S11.**

**Impacts of sampling and sequencing on diversity and population structure metrics.** Associations between number of individuals (A,B) and average number of sites per individual (C,D) in each population with diversity statistics. Relationships between principle component values and number of individuals within each population (E,F) as well as principle component values and mean coverage for each individual (G,H). Associations between ancestry proportions for cluster (K1-K3) and mean sequencing coverage for each individual (I-K). Mean coverage was calculated via QualiMap and ancestry proportions were calculated with NGSadmix for a representative run with K=3 (the most likely K value).

| Structural Variant | Fitness Measure | Genotype | | | Garden | | | Genotype:Garden | | |
|---|---|---|---|---|---|---|---|---|---|---|
| | | Chisq | df | p-value | Chisq | df | p-value | Chisq | df | p-value |
| HB7a1 | Survival (1st Year) | 0.1 | 1 | 0.751 | **60.15** | **3** | **5.46E-13** | 2.93 | 3 | 0.402 |
| HB7a2 | Survival (1st Year) | 0.56 | 1 | 0.454 | **50.92** | **3** | **5.08E-11** | **7.46** | **3** | **0.059** |
| HB7b | Survival (1st Year) | 0.08 | 1 | 0.773 | **58.86** | **3** | **1.03E-12** | 5.75 | 3 | 0.124 |
| HB9 | Survival (1st Year) | 0.28 | 1 | 0.595 | **44.06** | **3** | **1.47E-09** | 0.8 | 3 | 0.851 |
| HB13 | Survival (1st Year) | 0.05 | 1 | 0.825 | **30.43** | **3** | **1.12E-06** | 1.02 | 3 | 0.797 |
| HB7a1 | Survival (Flowering) | 2.32 | 1 | 0.128 | **70.53** | **3** | **3.28E-15** | 5.72 | 3 | 0.126 |
| HB7a2 | Survival (Flowering) | 0.01 | 1 | 0.944 | **60.33** | **3** | **5.00E-13** | 2.31 | 3 | 0.51 |
| HB7b | Survival (Flowering) | 0.77 | 1 | 0.38 | **64.63** | **3** | **6.03E-14** | 2.87 | 3 | 0.412 |
| HB9 | Survival (Flowering) | 0.16 | 1 | 0.692 | **53.15** | **3** | **1.70E-11** | 2.15 | 3 | 0.543 |
| HB13 | Survival (Flowering) | 0.07 | 1 | 0.787 | **35.15** | **3** | **1.13E-07** | 0.79 | 3 | 0.852 |
| | | F | df | p-value | F | df | p-value | F | df | p-value |
| HB7a1 | Total Seed Mass | No Variation in Sweden Garden | | | | | | | | |
| HB7a2 | Total Seed Mass | **7.41** | **1** | **0.007** | **7.08** | **3** | **1.54E-04** | 2.31 | 3 | 0.078 |
| HB7b | Total Seed Mass | 0.04 | 1 | 0.849 | **6.66** | **3** | **2.65E-04** | 0.47 | 3 | 0.703 |
| HB9 | Total Seed Mass | 0.77 | 1 | 0.382 | **7.04** | **3** | **1.63E-04** | 0.85 | 3 | 0.47 |
| HB13 | Total Seed Mass | 7.75 | 1 | 0.006 | **4.76** | **3** | **0.003** | 7.65 | **3** | **7.43E-05** |
| HB7a1 | Absolute Fitness | 0.32 | 1 | 0.57 | **32.29** | **3** | **2.20E-16** | **2.8** | **3** | **0.04** |
| HB7a2 | Absolute Fitness | 1.2 | 1 | 0.274 | **30.2** | **3** | **2.20E-16** | 1.86 | 3 | 0.135 |
| HB7b | Absolute Fitness | 1.14 | 1 | 0.286 | **31.29** | **3** | **2.20E-16** | **2.56** | **3** | **0.054** |
| HB9 | Absolute Fitness | 0.04 | 1 | 0.841 | **22.67** | **3** | **1.03E-13** | 0.35 | 3 | 0.79 |
| HB13 | Absolute Fitness | 1.33 | 1 | 0.249 | **6.68** | **3** | **2.03E-04** | **9.59** | **3** | **3.74E-06** |

**Table S1.**
**Associations between fitness variables and haploblock genotypes across four common gardens**. ANOVA results based on a type-III sum-of-squares. Both survival measures were modeled within a generalized linear model with a binomial distribution and logit link function. Total Seed Mass does not include individuals that did not survive to flowering. Absolute fitness is measured as total seed mass with individuals not surviving to flowering having zero total seed mass. Total seed mass and relative fitness were log+1 transformed to improve model fit.

**Data S1. (separate file)**
Population summary statistics from population genomic analyses.

**Data S2. (separate file)**
Gene ontology analysis for each haploblock.

**Data S3. (separate file)**
Common-garden summary statistics for each haploblock.

**Data S4. (separate file)**
Fitness-associated loci from the common garden study.

**Data S5. (separate file)**
Differentially Expressed Genes from the RNAseq study.
